# Characterization and analysis of the transcriptome in *Arapaima gigas* using multi-tissue RNA-sequencing

**DOI:** 10.1101/2020.09.29.317222

**Authors:** Danilo L. Martins, Leonardo R. S. Campos, André M. Ribeiro-dos-Santos, Ana Carolina M. F. Coelho, Renata L. Dantas, Pitágoras A. A. Sobrinho, Tetsu Sakamoto, Amanda F. Vidal, Glória T. Vinasco-Sandoval, Paulo P. Assumpção, Ândrea K. C. R. Santos, Rodrigo J. S. Dalmolin, Sandro J. de Souza, Sidney Santos, Jorge E. S. de Souza

**Author notes:** Correspondence: Dr. Jorge E. S. de Souza.

## Abstract

*Arapaima gigas* is a giant bony tongue air-breathing fish, and a promising species for aquaculture due to its particular features. However, there is still a lack of information on its biology and few transcriptome studies are available. Our aim was to characterize the transcriptome of arapaima in order to shed light on molecular networks contributing to its unique traits. Through RNA-sequencing, we generated a transcriptome from eight tissues (brain, pituitary, heart, muscle, kidney, lung, ovary, and testis) collected from arapaima adults specimens. Using a genome-guided strategy associated with homologous protein evidence, 57,706 transcripts were assembled, which aligned to 23,353 high confidence protein-coding genes. The analysis revealed a global view of expression patterns, as well as it allowed us to identify tissue-specific gene clusters, transcription factors within the clusters, and to compare expression patterns between male and female. These analyses has generated tissue-specific and sex-biased transcriptome profiles, which will be helpful to understand its molecular biology, evolution, and also guide future functional studies of the arapaima.

## Introduction

*Arapaima gigas*, also known as Pirarucu or Paiche, is one of the largest freshwater fishes in South America, whose adults body weight can reach up to 250 kg [1] and a length of up to three meters [2] [3]. It belongs to the bonytongues (order Osteoglossiformes) [2] [4] and it is naturally distributed in Amazonas and Tocantins-Araguaia basins [1] [3] [4], and it has also been introduced into the Bolivian Amazon [3] [5]. This emblematic fish has been an extensive economic resource for communities and is an attractive candidate for aquaculture development [1] [2]. It exhibits the fastest growth rates among Amazonian cultivated fishes, reaching 10-15 kg/year [6]. In the wild, this air-breather fish lives preferentially in lentic environments [7], tolerates low oxygen levels dissolved in the water and is less susceptible to ammonia or nitrite intoxication [2]. The reproduction behavior is complex, which includes breeding pairs, nest building and parental care, and significant external sexual dimorphism is not displayed [1] [3]. Methods usually used for sexual identification include biopsy, ultrasonography and endoscopy [8]. However, sexual maturity is reached only after 3 to 5 years of age and only the left gonad is functional both in males and females, which makes biopsies difficult and labor intensive to perform [2].

Recently, due to the rapid development of next-generation sequencing technologies, genome and transcriptome analysis become powerful tools for identifying candidate genes [9] related to these important traits, and it has provided an advancement in availability of arapaima genomics and transcriptomics resources [3]. The ability to elucidate the basis of complex traits has been dramatically improved by the availability of reference genomes for *A. gigas* [2] [4]. To date, their major impact has been through enabling evolutionary analysis and application of analysis of gene family dynamics. The availability of the *de novo* transcriptome [3] has also enabled performing differential gene expression analysis. Unfortunately, there is still a lack of knowledge about molecular and biochemical mechanisms involved in its fast growth, metabolism and sexual development, which interfered in the widespread establishment of *A. gigas* in aquaculture [2]. Future genomic and transcriptomic works providing detailed knowledge of organismal biology would greatly provide useful information for establishing this remarkable fish in aquaculture.

Here, a functionally annotated *A. gigas* transcriptome generated from multiple tissues (brain, pituitary, lung, kidney, heart, muscle, ovary, and testis) of two adults is presented. The genome-guided transcriptome assembled served as a reference for read mapping and estimating gene expression. Because using multiple tissues, specific genes were identified to better characterize functionally each tissue. Additionally, transcription factors potentially involved in tissue-specific gene expression patterns were also identified. Sex-biased genes were further investigated in order to identify potential candidates related to sexual determination. Collectively, these analyses shed light on important questions in organismal biology and complex traits, showcasing this transcriptome as a valuable approach to gain insights into molecular mechanisms underlying specific traits of this remarkable fish.

## Results

### Transcriptome sequencing, assembly and annotation

The overall process of transcriptome sequencing, assembly, annotation and subsequent analyses is summarized in Figure 1. The Illumina NextSeq 500 sequencing produced a total of 358,617,594 raw reads across eight organs (brain, lung, pituitary, kidney, muscle, heart, ovary and testis) of *Arapaima gigas*. The raw transcriptome sequences have been submitted to the NCBI database and are accessible under Accession no. **PRJNA665688**. After trimming and rRNA removal, a total of 260,680,050 (∼73%) high-quality reads were retained for transcriptome assembly.

**Figure 1.**
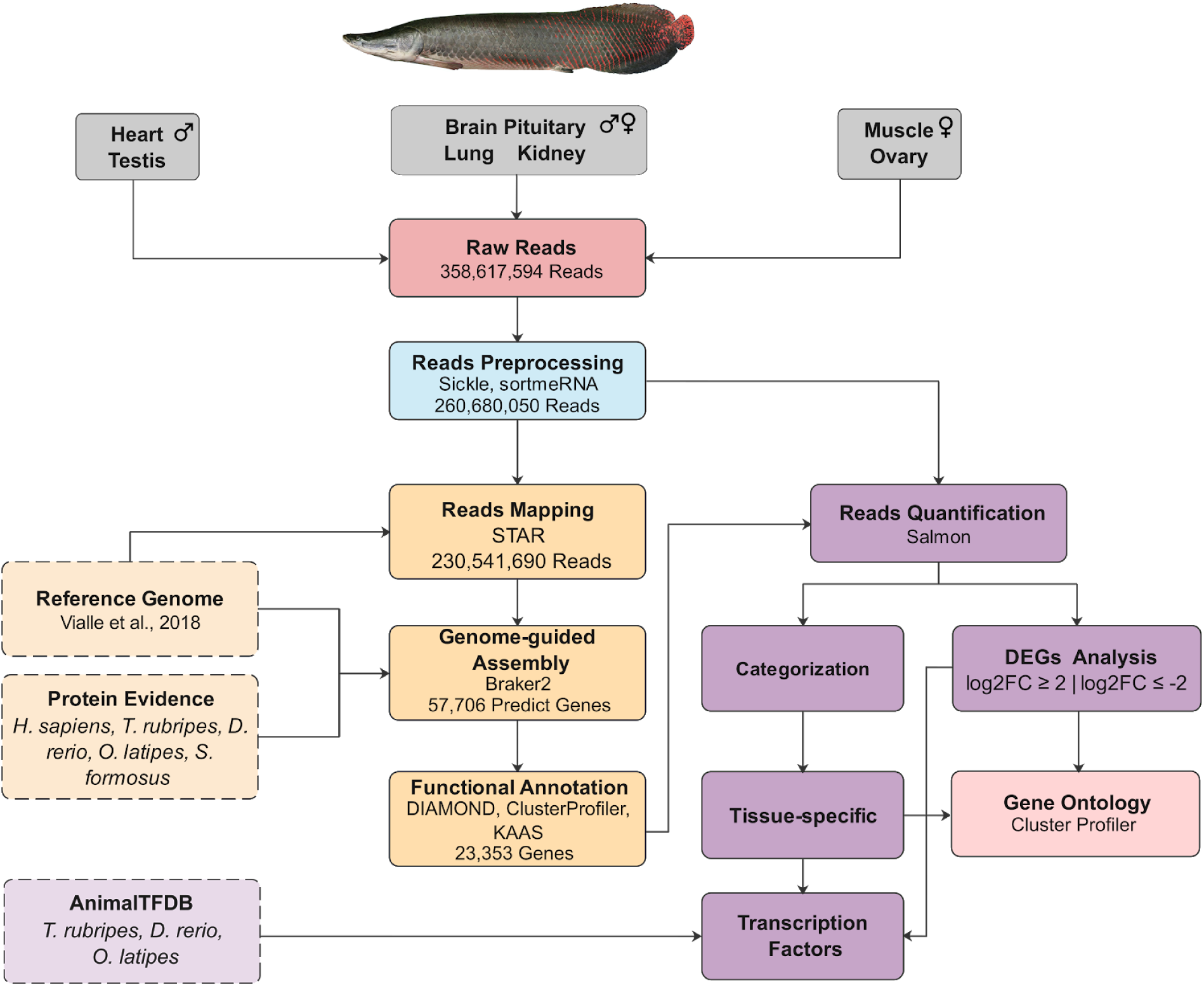
Overall workflow of bioinformatics analyses. Sequential workflow steps from the initial raw input data to data processing analyzes.

Out of these retained reads, 230,541,690 were mapped against the genome reference and submitted to Braker2 pipeline to genome-guided gene prediction (Table 1). A total of 57,706 transcripts sequences were assembled, ranging from 21 to 31,680 bp, and the average length of 1,156 bp (Fig. 2). Sequences less than 180 bp were removed and finally, a total of 57,244 sequences were submitted for annotation. Out of these transcripts sequences, 35,216 showed significant alignment against the *S. formosus* and NCBI non-redundant protein sequences database (BLASTP within DIAMOND, e-value 1e-5, keeping top hit).

**Table 1.** Summary information for the Arapaima gigas transcriptome from the all 12 libraries generated separately.

| Tissues | Raw Reads | Trimmed Reads | Ribosomal RNA | Filtered Reads | Mapped Reads | % Mapped reads |
| --- | --- | --- | --- | --- | --- | --- |
| Brain Female | 34,238,962 | 27,222,140 | 924,418 | 26,297,722 | 23,939,138 | 91.03% |
| Brain Male | 27,003,784 | 23,800,768 | 1,623,032 | 22,177,736 | 20,713,252 | 93.40% |
| Pituitary Female | 25,252,632 | 20,925,752 | 1,003,100 | 19,922,652 | 18,297,270 | 91.84% |
| Pituitary Male | 19,560,392 | 14,157,718 | 1,201,786 | 12,955,932 | 11,128,788 | 85.90% |
| Kidney Female | 23,939,386 | 17,063,434 | 840,114 | 16,223,320 | 13,821,650 | 85.20% |
| Kidney Male | 27,014,308 | 17,761,092 | 1,536,756 | 16,224,336 | 13,547,540 | 83.50% |
| Lung Female | 18,493,824 | 13,684,668 | 1,072,262 | 12,612,406 | 10,881,630 | 86.28% |
| Lung Male | 42,443,468 | 32,457,950 | 1,741,642 | 30,716,308 | 26,849,952 | 87.41% |
| Heart Male | 20,023,278 | 14,117,204 | 1,583,620 | 12,533,584 | 10,396,914 | 82.95% |
| Muscle Female | 33,566,070 | 23,119,982 | 940,804 | 22,179,178 | 18,933,380 | 85.37% |
| Ovary | 68,657,938 | 59,223,394 | 1,204,302 | 58,019,092 | 53,206,006 | 91.70% |
| Testis | 18,423,552 | 12,655,458 | 1,837,674 | 10,817,784 | 8,826,170 | 81.59% |

**Figure 2.**
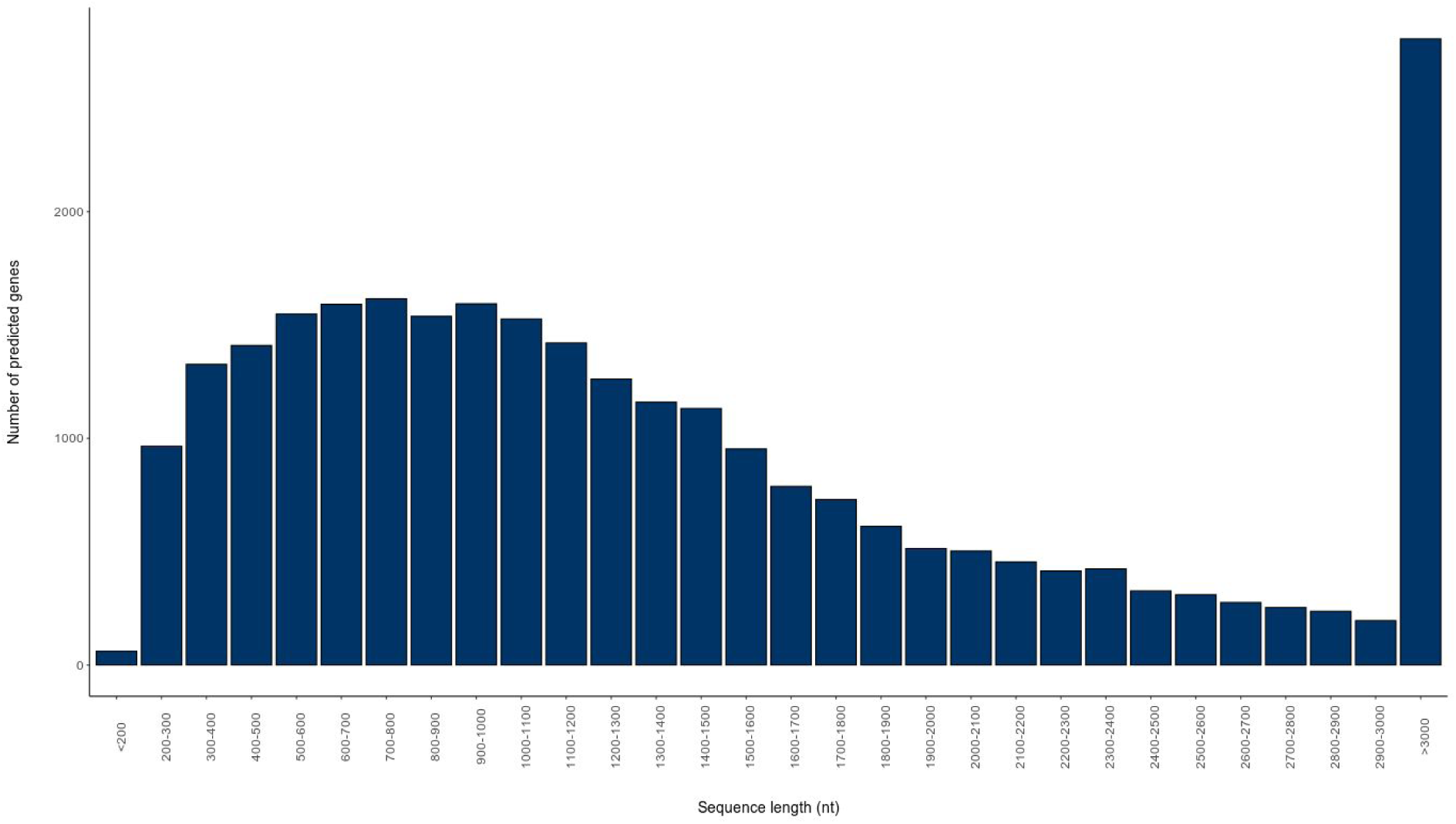
Length distribution of assembled transcripts from different tissues of *A. gigas*. The x-axis represents the sequence length and the y-axis represents the number of assembled transcripts.

### Global gene expression patterns and assessment of tissue-specific genes

A total of 23,353 transcripts, including isoforms, were identified in the transcriptome and quantified using Salmon to evaluate the transcriptional activity of the genes. Each transcript was then categorized according to expression levels (Supplementary Fig. S1). Overall, approximately 48% (11,108) of the genes were moderated or low expressed across all analyzed tissues, 19% (4,457) showed mixed expression, 11% (2,674) were enriched in a group of tissues, and ∼4% (1,026) did not show significant expression in any of the tissues. Only ∼17% (3,886) were enriched in one tissue and ∼1% (202) of the genes were highly expressed in one analyzed tissue. Transcripts highly expressed or enriched in one analyzed tissue are considered hereafter as tissue-specific genes. The relative expression of identified tissue-specific transcripts is shown in Figure 3.

**Figure 3.**
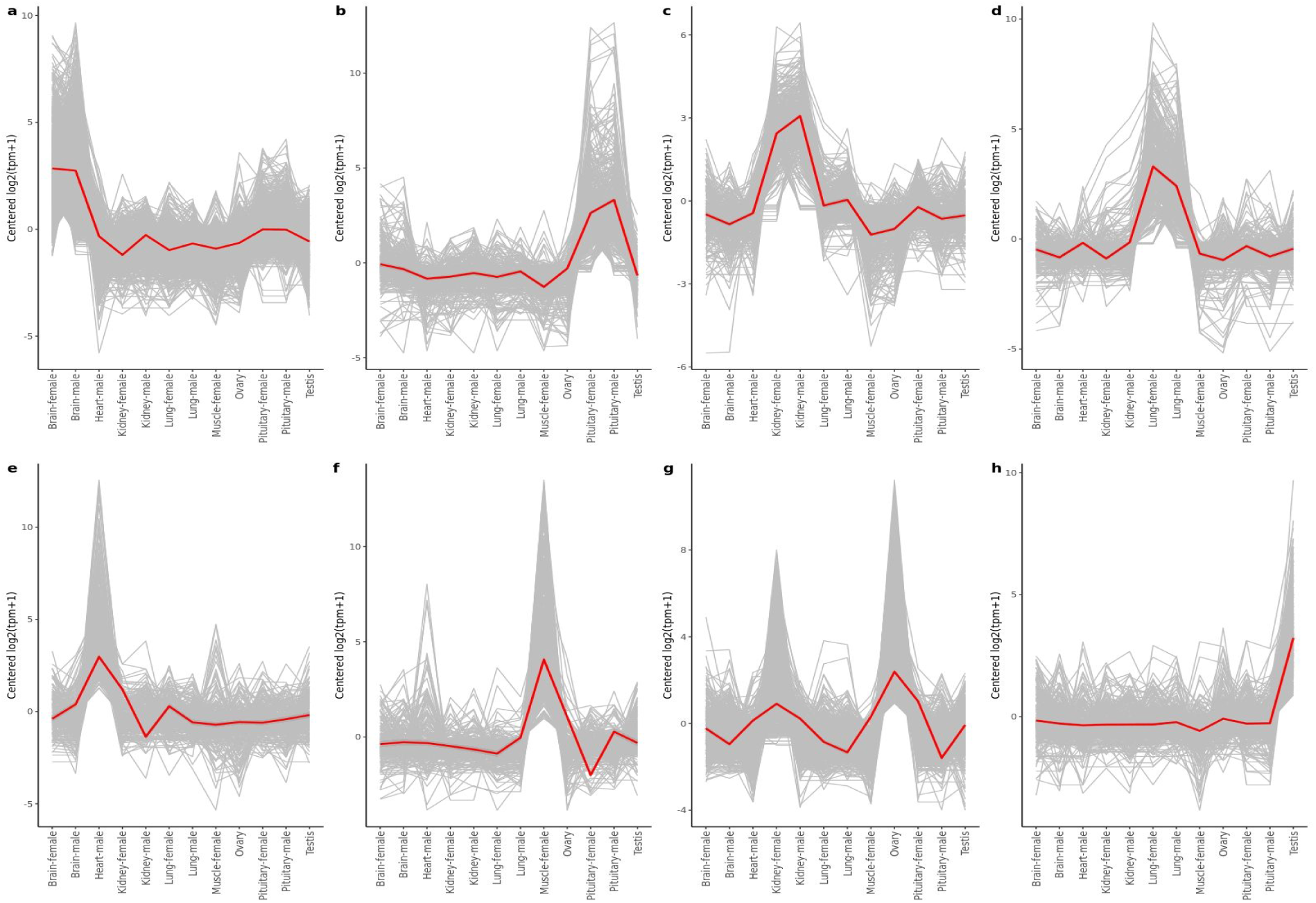
Relative expression of tissue specific genes across tissues. The y-axis of each graph represents the mean-centered log2 (TPM+1) value. Expression levels of each tissue-specific genes is shown in gray, and the mean expression of them in **(a)** brain, **(b)** pituitary, **(c)** kidney, **(d)** lung, **(e)** heart, **(f)** muscle, **(g)** ovary, and **(h)** testis is shown in blue.

The clustering analysis of tissue-specific genes revealed distinctive expression profiles for each tissue. In addition, it also indicates similar expression profiles between both sexes. This finding was corroborated in a comparative analysis with others transcriptome public data (PD) available (NCBI BioProject PRJNA353913) (Fig. 4). In the broader context, specific tissue data has remained consistent either showing a grouping of our analysis with that of the PD for the tissue pairs in common (ovary and testis) and at the same time a more defined clustering in the tissues that we don’t have data (skin and liver) (Supplementary Fig. S2). Tissue-by-tissue comparison showed that brain (1,423), ovary (647), and testis (470) had the largest number of tissue-specific genes, followed by pituitary (357), kidney (342), and lung (315), with the lowest number in muscle (268) and heart (266). An UpSet plot was constructed to show the presence/absence of genes across tissues (Fig. 5). The most abundant tissue-specific gene were *mbpa* in brain, hypothetical protein Z043_117078 (KPP64562.1) in kidney, gonadotropin alpha subunit in pituitary, hemoglobin subunits alpha- and beta-like (*LOC108935728, LOC108935782*) in lung, *ckmb* in muscle, *LOC108922717* in heart, *LOC114911227* in ovary, and *hp* in testis.

**Figure 4.**
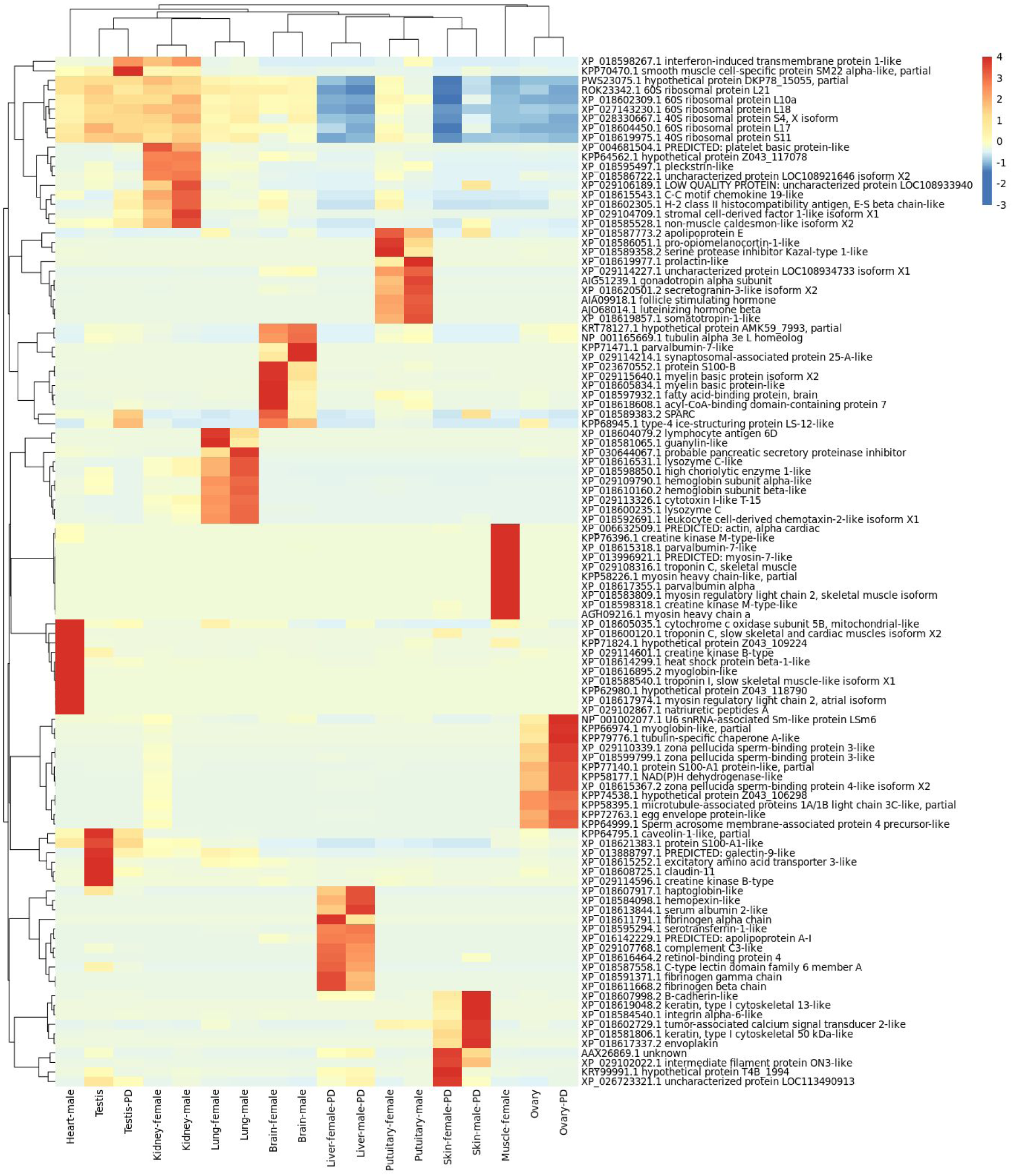
Top 10 tissue-specific gene expressions. Heatmap and dendrograms of the 10 most highly expressed genes in each tissue transcriptome from this work and from a public database (PD). The color gradient indicates the relative expression levels in TPM. Highly expressed genes are plotted in red, while low expressed genes are plotted in blue.

**Figure 5.**
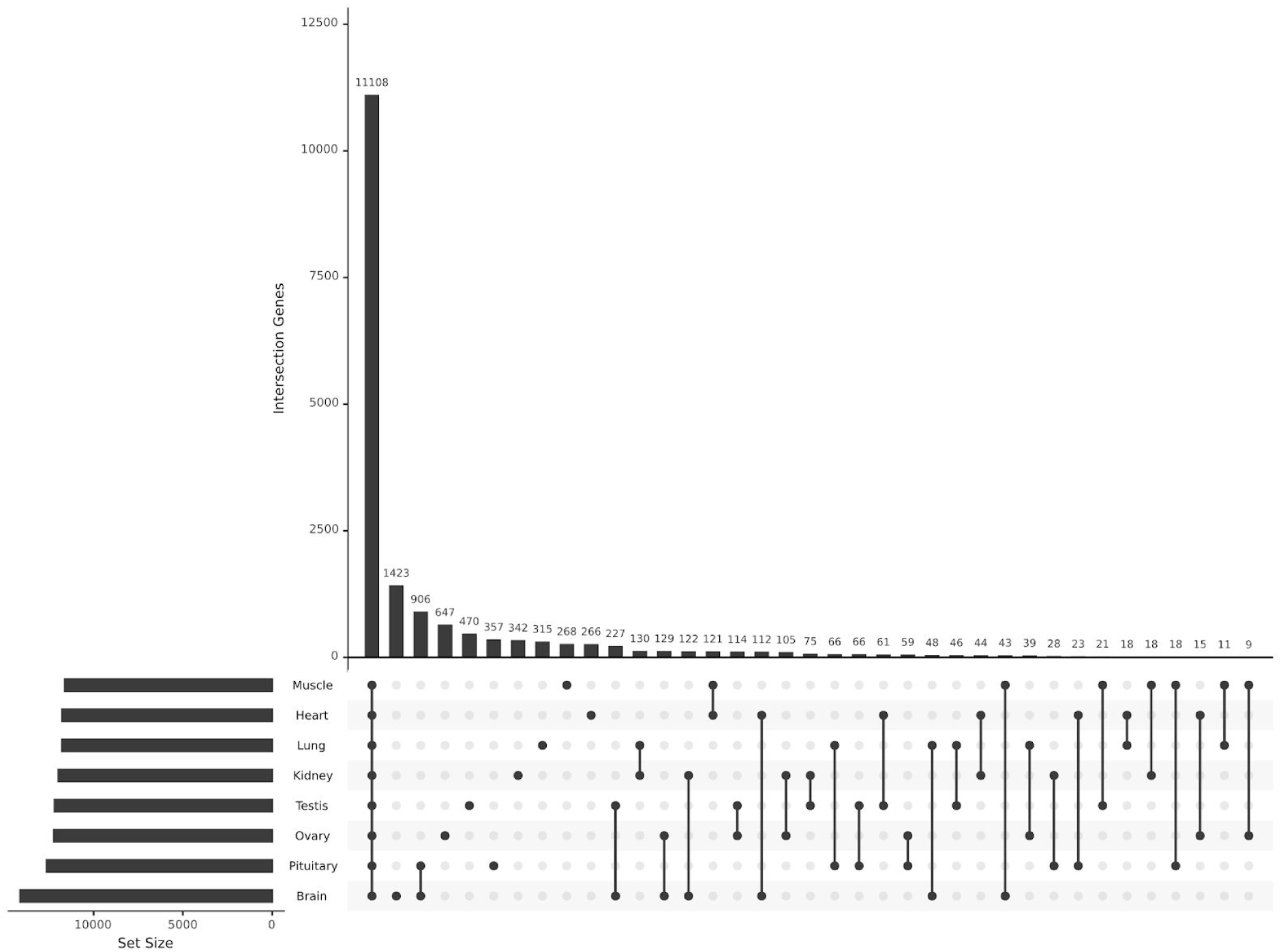
Distribution of specific and shared transcripts across tissues. UpSet plot showing the number of specific and shared transcripts found in each tissue. The bar chart on the left indicates the total number of transcripts for each tissue. The upper bar chart indicates the number of transcripts found in one tissue or the intersection between sets of transcripts found in two or more tissues. Connected dots on the bottom panel indicate which tissues are being considered to determine the intersection.

Tissue-specific genes were enriched for GO terms, covering all three domains: BP, MF and CC. Focusing on BP categories, GO enrichment analyses indicated that synaptic signaling (GO:0099536) was the most significant term in brain, cell cycle process (GO:0022402) in ovary, embryonic organ morphogenesis (GO:0048562) in testis, head development (GO:0060322) in pituitary, immune response (GO:0006955) in kidney, anion transport (GO:0006820) in lung, and muscle structure development (GO:0061061) in both muscle and heart. Top 10 GO terms of each tissue cluster is shown in Supplementary Figure S3.

A pathway enrichment analysis was also performed to further investigate the biological functions of tissue-specific genes. All these genes were mapped to reference pathways in the KEGG database to identify the metabolic pathways in which these genes may be involved (Table 2). The calcium signaling pathway was the most significantly enriched pathway for brain-, kidney- and muscle-specific genes, followed by MAPK signaling pathway, regulation of actin cytoskeleton and cardiac muscle contraction, respectively. Melanogenesis was the most enriched pathway for pituitary-, whereas glyoxylate and dicarboxylate metabolism pathway was the most enriched pathway lung-, WNT signaling pathway for heart-, cell cycle for ovary-, and cytokine-cytokine receptor interactor pathway for testis-specific genes.

**Table 2.** Top 2 most significant (p-value ≤ 0.05) KEGG pathways for tissue-specific genes.

| ID | Description | p-value | Count | Tissue |
| --- | --- | --- | --- | --- |
| ko04020 | Calcium signaling pathway | 0.00027 | 18 | Brain |
| ko04310 | Wnt signaling pathway | 0.00308 | 12 | Brain |
| ko04916 | Melanogenesis | 0.00016 | 6 | Pituitary |
| ko04744 | Phototransduction | 0.00016 | 4 | Pituitary |
| ko00630 | Glyoxylate and dicarboxylate metabolism | 0.00077 | 3 | Lung |
| ko00260 | Glycine, serine and threonine metabolism | 0.00155 | 3 | Lung |
| ko04020 | Calcium signaling pathway | 0.00001 | 8 | Kidney |
| ko04810 | Regulation of actin cytoskeleton | 0.00803 | 5 | Kidney |
| ko04310 | Wnt signaling pathway | 0.00041 | 6 | Heart |
| ko04261 | Adrenergic signaling in cardiomyocytes | 0.00499 | 5 | Heart |
| ko04020 | Calcium signaling pathway | 0.00037 | 8 | Muscle |
| ko04260 | Cardiac muscle contraction | 0.00015 | 6 | Muscle |
| ko04110 | Cell cycle | 0.00001 | 11 | Ovary |
| ko04114 | Oocyte meiosis | 0.00001 | 10 | Ovary |
| ko04060 | Cytokine-cytokine receptor interaction | 0.04379 | 4 | Testis |
| ko04512 | ECM-receptor interaction | 0.02721 | 3 | Testis |

### Expressed TFs within tissue-specific genes

To further characterize regulatory mechanisms, a list of tissue-specific genes involved in regulation of gene expression was generated using the AnimalTFDB database (Supplementary Fig. S4). A total of 148 transcription factors were identified among brain-specific transcripts including *id4, tsc22d1*, zinc fingers (*zic2a, zic1, LOC108920431*), and homeobox proteins (*en2a, LOC108922262)*. Additionally, 68 and 50 transcription factors were identified in ovary- and testis-specific transcripts, respectively. The most highly expressed transcription factor in ovary was *si:dkeyp-75b4*.*8*, followed by *LOC115203203, si:dkey-208k4*.*2, figla* and *irx7*. While highly expressed TFs in testis included *si:ch211-14a17*.*10, LOC114785964, bnc1*, and *wt1*. Furthermore, 50 TFs were also identified in pituitary including *xbp1, ascl1a, six6a, lmo4*, and insulinoma-associated proteins (*insm1a, insm1b*). A total of 36 TFs were identified in both muscle- and kidney-specific transcripts. Highly expressed TFs identified in muscle included *casq1a, myod1* and myosin heavy chains (*LOC114909085, LOC115813863, LOC114910488*). While in kidney the list included *plek, tlx, nkx2*.*7*, and interferon regulatory factors (*irf8* and *LOC108937748*). Within heart- and lung-specific genes, 33 and 24 TFs were identified, respectively. In the heart, *desmb, LOC108930075*, and *hand2* were among the highest expressed TFs. In lung, identified highly expressed TFs included *LOC108931228, foxi3b, cebp1, zgc:92380*.

### Sex differences in gene expression

Putative sex-biased genes were identified based on their abundance levels in samples from different sexes. A total of 383 genes were identified as female-biased, most of them highly expressed in ovary and brain, while 224 putative male-biased genes were identified, especially in brain and testis. The remaining transcripts were considered unbiased.

Among the most abundant putative female-biased genes there were zona pellucida sperm-binding proteins (*LOC114911227, LOC108929618, LOC108938882*). Other notable female-biased genes included *Figla*, and *Gdf9* transcripts were also overexpressed in the ovary. While the most abundant male-biased genes include *hp, rrad, cldn11, anxa2*, overexpressed in testis. Another notable male-biased gene is *dmrta1*, which is overexpressed in the male pituitary. The top 10 sex-biased genes list is included as Supplementary Table S1.

To interpret the global transcript function, GO classification revealed both female- and male-biased genes assigned to functional groups of the three ontologies. Enriched GO terms identified in putative female-biased and male-biased genes demonstrated similar patterns (Fig. 6). Among the highest enriched biological process terms were cellular process (GO:0009987), biological regulation (GO:0065007) and metabolic process (GO:0008152). The terms reproduction (GO:0000003) and reproductive process (GO:0022414) appeared exclusively for the female-biased genes. In the CC category, cell (GO:0005623), cell part (GO:0044464) were the most abundant groups in both female and male-biased groups. The term membrane-enclosed lumen (GO:0031974) also appeared only for female-biased genes. Within the MF category, binding (GO:0005488), catalytic activity (GO:0003824) and transporter activity (GO:0005215) were the top three functional terms. In addition, sex-biased genes were further characterized by KEGG database. Metabolic pathways with greatest representation included oocyte meiosis, cell cycle, glutathione metabolism and progesterone-mediated oocyte maturation for female-biased genes, and calcium signaling pathway, endocytosis and apelin signaling pathway for male-biased genes. The most significant sex-biased genes pathways is included as Supplementary Table S2.

**Figure 6.**
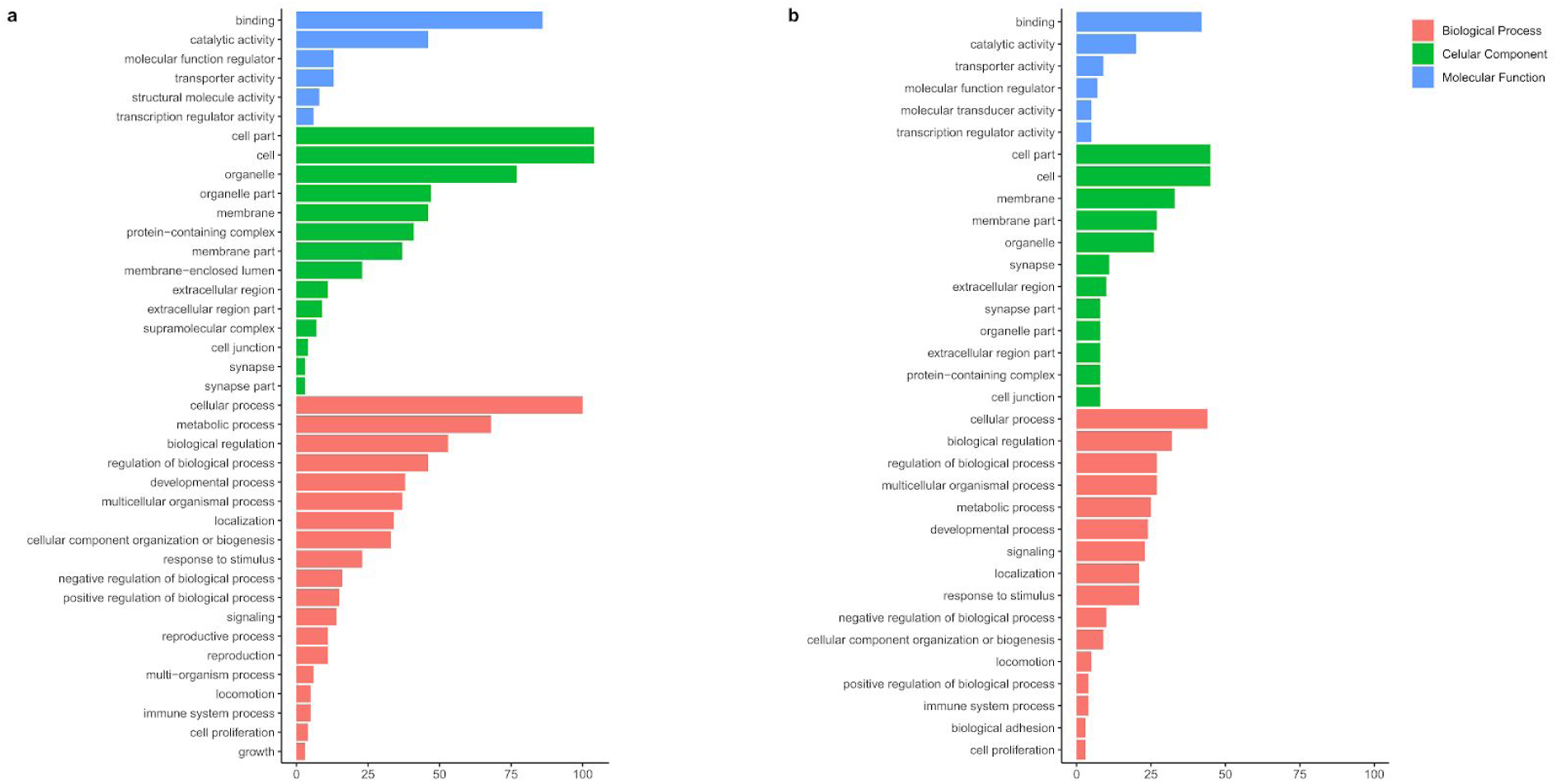
GO terms distribution of sex-biased identified genes. (**a**) Distribution of GO terms from transcripts identified mostly expressed in female tissues. (**b**) Distribution of GO terms from male-biased transcripts. Bar plots describe the distribution of cellular components (green), molecular function (blue) and biological processes (salmon) categories. The size of each bar is proportional to the number of counts classified in each functional category.

### Expressed TFs within sex-biased genes

Transcription factors that may be regulating the gene expression of the sex-biased genes were also evaluated using the AnimalTFDB database (Supplementary Fig. S5). A total of 30 transcription factors were identified among female-biased genes, including *Figla, si:dkeyp-75b4*.*8, cth1, zglp1, LOC115202771* and *LOC115203203*. These transcription factors had at least 50-fold higher expression compared to male-biased genes. Within male-biased genes, 25 transcription factors were found, including zinc fingers (*zic1, zic2a, LOC108927773, LOC108920431*), *LOC108925283*, and *LOC108932644*. These transcription factors had at least 50-fold higher expression compared to female-biased genes. In addition, *neurod2* and *LOC108922262* were also identified as male-biased with expression higher than 30-fold.

## Discussion

Transcriptome sequencing has been proposed as an efficient method for gene discovery and analysis of differential gene expression in non-model organisms. Thus availability of transcriptomic data from multiple tissues can elucidate important questions in evolutionary and organismal biology and serve as a platform to guide future functional genomic studies [10]. In the present study, RNA-Seq from multiple tissues was used to generate a comprehensive transcriptome using genome-guided gene prediction strategy in combination with external protein evidence. Here, the *A. gigas* transcriptome assembly provides a potential resource for further understanding molecular mechanisms underlying the remarkable traits of this remarkable fish. The number of annotated transcripts is still low due to the phylogenetic distance of arapaima and consequently limited availability of properly annotated genomes of orthologous species. However, our data suggest our transcriptome from one male and one female (8 tissues each) has good coverage. This transcriptome assembly is valuable for molecular and genetic studies of *A. gigas*, and the closely related species. Future studies would ideally have more replicates and more tissues from both sexes, and juveniles specimens.

In comparison, the number of annotated transcripts (23,353) and the mean transcript size (1,156 bp) were slightly higher to those recorded in a previous *A. gigas* transcriptome study (20,219, 717.95 bp) assembled by using a *de novo* strategy [3]. These differences may be due to the difference of assembly strategies and algorithms, sequencing platforms and the number of tissues sequenced to assemble the reference transcriptome.

Identifying the genes expressed in a given tissue represents important baseline information essential for understanding tissue function and physiology more broadly [11] in multicellular organisms. In our data, tissue-specific expressed transcripts and transcription factors generally agreed with expectation about the specific role of each organ. For example, muscle-specific genes and transcription factors were associated with muscle development and muscle contraction, whereas heart-expressed genes and TFs were associated with cardiac muscle tissue development, blood circulation and heart contraction, which aligns with functions of the muscle and heart, respectively. The brain had the most tissue-specific genes and transcription factors, which were associated with synaptic signaling, neurotransmitter transport, and other neural processes.

In addition, the ovary expressed the second highest number of tissue-specific genes and showed the most significant overrepresentation for terms pertaining to cell cycle, regulation of cell cycle, likely having to do with oogenesis events, and reproduction process. In fish ovaries, mitotic germ cells are histologically observed as oogonia and thus the proliferation of oogonia can supply mature eggs in mature ovaries [12]. Furthermore, as arapaima are partial spawners, they never have an empty ovary [13]. In testis, specific genes and transcription factors were associated with terms related to testis-specific functions, such as reproduction and motility.

Furthermore, genes with maximum expression in the kidney were mainly associated with immune response. This finding is consistent with the observation of major functions of this tissue. The teleost kidney contains two sections: the head kidney, which comprises the anterior portion, and the trunk kidney which extends along the dorsal wall of the body cavity [14]. The head kidney is an important endocrine and hematopoietic-lymphoid organ [15], which plays a key role in initiation of the immune response [16]. The lung showed a large number of specific transcripts largely related to transmembrane ion transport, which aligns with the function of this organ. The respiratory gas bladder, analogous to a lung and here referred as such, of the adult arapaima is used to exchange gases with its environment [13]. As an obligate air-breather, the arapaima has an extremely thin air–blood barrier in the swim bladder, which enables the O_2_ uptake and CO_2_ excretion [17].

Ultimately, the pituitary had more general overrepresented GO terms. However, genes with highest expression in this tissue had relatively high tissue-specificity, which were highly related to endocrine hormones. For example, *pomca, gonadotropin alpha subunit* (AIG51239.1), *luteinizing hormone beta* (AJO68014.1), and *gh1*. The *pomca* gene is expressed in both the anterior and intermediate lobes of the pituitary gland [11]. Similarly, gonadotropin, somatotropin and luteinizing hormones are produced by the pituitary gland. The anterior lobe contains ACTH, gonadotropin, prolactin, and growth hormone secreting cells, while the intermediate lobe contains cells that can produce melanotropin, adrenocorticotropin and somatolactin [13].

To investigate sex-specific gene expression patterns in these tissues, we analyzed and compared differences between male and female transcriptomes. A global view of sex-biased expression in these tissues was obtained. We observed that sex-biased genes were preferentially expressed in gonads and brain, the most important tissues in sexual development and reproduction [18]. Although molecular mechanisms of sex determination in *A. gigas* remains unknown, the identification of sex-biased transcripts is a powerful tool for the identification of potential candidates implicated in sex differentiation and sex maintenance in this species.

A closer look into sex-biased genes that were up-regulated in female tissues, revealed numerous candidates with a notable role in sex differentiation in other species, including teleosts. For example, the *Figla* gene appears as female-biased with high expression in the ovary. This folliculogenesis transcription factor mediates several development process, including fertilization and early embryogenesis [19], and up-regulates zona pellucida (zp) genes which encodes products used to build the vitelline envelope surrounding eggs in teleosts [18]. Similarly, genes encoding several zp proteins were also up-regulated in female tissues. ZP are glycoproteins of the fish chorion [20], and appeared to be expressed in ovary in humans, mice, and some teleosts [18]. The same expression pattern was shown for *Gdf9*, an oocyte-specific growth factor, which is required for follicular development and follicle growth in different vertebrates, including mammals, chicken, and zebrafish [21].

Contrary to other transcriptomic studies in teleosts [22] [23], our findings have not pointed out a male-biased transcripts over-expression tendency, probably due to the criterias used on the sex-biased classification. A number of male-biased genes were identified, but with low levels of TPM. Nevertheless, the *dmrt1a* gene appeared up-regulated in the male pituitary. The *dmrt* genes have attracted considerable interest because of their involvement in sex determination and differentiation across invertebrate and vertebrate species [24], and the role of the autosomal *dmrt1a* in testis maintenance is well established in medaka fish [25].

In conclusion, this study allowed us to assemble a comprehensive transcriptome of *Arapaima gigas*. A collection of expressed genes in multiple arapaima tissues was cataloged. A number of tissue-specific genes clusters were identified, and their functional analyses enable further characterization of each tissue. Moreover, highly-expressed transcription factors (TFs) in each tissue-specific gene cluster were also identified. Ultimately, our findings have demonstrated gene expression differences between male and female adults of arapaima. Functional pathways behind sex-biased expression revealed the recruitment of known sex-related genes either to male or female in teleosts. Taken together, these analyses provide valuable information for understanding the molecular mechanisms underlying the arapaima unique traits and will be helpful to perform future functional studies of the arapaima.

## Methods

### Tissue sampling and ethics statement

Two adults *Arapaima gigas* specimens (one male and one female) were captured to sample collection. The male was dissected to obtain samples of brain, lung, pituitary, kidney, heart and testis while the female was dissected to obtain samples of brain, lung, pituitary, kidney, muscle and ovary. Samples were stored immediately in RNAIater (Thermo Fisher Scientific, Waltham, MA) stabilization solution at -20 °C until RNA extraction. All sampling procedures were conducted in strict accordance with the standards of the Federal University of Pará animal protocol. All efforts were made to minimize suffering of the animals.

### RNA extraction and sequencing

Total RNA was extracted from tissue samples using TRIzol reagents (Thermo Fisher Scientific, Waltham, MA) according to instructions of the manufacturer. Purified RNA integrity, quality and concentration were determined using Qubit 2.0 Fluorometer (Invitrogen, CA, USA) and Agilent 2200 TapeStation System (Agilent Technologies, Santa Clara, CA, USA). The RNA with good integrity was utilized to construct the library. The cDNA library was generated from 1 µg of total RNA using TruSeq RNA Sample Prep Kit (Illumina, San Diego, CA, USA) following manufacturer’s protocols. Paired-end sequencing was performed on the Illumina NextSeq 500 platform using NextSeq 500/550 High Output kit (Illumina, San Diego, CA, USA).

### Data processing and transcriptome assembly

Raw reads quality was assessed using FastQC tool [26] and low quality reads and adapters were then discarded by using Sickle v1.33 [27] under default parameters. RNA ribosomal contamination was removed by using <u>SortMeRNA</u> v2.1 [28]. After removal of low quality reads, clean reads were retained and mapped to the first *Arapaima gigas* draft genome (NCBI:txid113544, FishBase ID: 2076) [4], hereafter referred to as genome reference, using STAR aligner v.2.6.0 software [29] with default settings. Resultant aligned reads were provided for training GeneMark-ET [30] and AUGUSTUS [31] within BRAKER2 pipeline v.2.1.5 [32] [33] [34] along with the *A. gigas* genome reference to perform genome-guided gene prediction. Protein sequences from following Ensembl species: *Homo sapiens* (human), *Takifugu rubripes* (Japanese puffer), *Danio rerio* (zebrafish), *Oryzias latipes* (Japanese medaka) and from *Scleropages formosus* (Asian arowana), the closest relative to *A. gigas*, were also submitted as extrinsic evidence to training AUGUSTUS with GenomeThreader [35].

### Functional annotation

To provide comprehensive functional annotation of the final unigene set, predicted sequences from genome-guided assembly were searched against the *S. formosus* protein database (NCBI:txid113540, Accession no. GCF_900964775.1) using BLASTP within DIAMOND software v.0.9.19 [36], with an e-value cutoff of 1e-5, identity of ≥ 75%. Only the best hit for each transcript was reported. Transcripts sequences were also compared to the NCBI non-redundant database (nr) (e-value cutoff 1e-5, best hit chosen). The R package ‘mygene’ [37] was used to assign gene symbols from NCBI RefSeq to predicted transcripts.

### Quantification of expression level and categorization of expression profiles

Predicted transcripts expression levels were evaluated using Salmon 0.14.1 [38] as transcripts per million (TPM). Reads from all sample tissues (kidney, lung, pituitary, brain, muscle, ovary, testis and heart) were mapped independently against the respective assembled transcriptome. Each predicted transcript was then categorized according expression levels into (a) ‘highly expressed’ (50-fold higher expression in one tissue compared to all other tissues), (b) ‘tissue enriched’ (5-fold higher expression level in one tissue compared to all other tissues), (c) ‘group enriched’ (5-fold higher average expression levels in a group of two or more tissues compared to all other tissues), (d) ‘mixed high’ (higher than 10 TPM expression level in up to the seven tissues), (e) ‘mixed low’ (expressed in up to seven tissues and less than 10 TPM expression level in at least one of these tissues), (f) ‘moderate expressed’ (higher than 10 TPM in all tissues), (g) ‘low expressed’ (expressed in all tissues and less than 10 TPM expression level in at least one of these tissues) and (h) ‘not significative’ (less than 1 TPM in all tissues).

### Identification of sex-biased differentially expressed genes

To identify differentially expressed genes between male and female, the expression fold change was calculated. Predicted transcripts from muscle or heart samples were eliminated from this analysis in order to avoid noise results due the fact that there were no paired samples. Transcripts were considered significantly differentially expressed between sexes when log2FC ≥ 2 (Male > Female) or log2FC ≤ -2 (Female > Male) and gene expression ≥ 100 TPM in at least one tissue. These stringent criteria were selected owing to the fact that there were no replicates.

### Identification of transcription factors (TFs)

In order to determine transcription factors (TFs) within genes categorized as highly expressed or tissue enriched (tissue-specific), and differentially expressed sex-biased genes, BLASTP within DIAMOND software (v.0.9.19) was performed (e-value 1e-5) against the transcription factor databases for *T. rubripes, D. rerio* and *O. latipes* downloaded from the AnimalTFDB database [39].

### Functional enrichment of tissue-specific and sex-biased genes

To infer functions of classified tissue-specific and sex-biased genes, Gene Ontology (GO) functional-enrichment analyses at level 2 were performed using the R package ‘clusterProfiler’ [40]. Genome of *D. rerio* (zebrafish) was used as the reference gene list due to the lack of other closest species in the available databases. Results were categorized as biological process, molecular function, and cellular components. The predicted sequences were also submitted to KEGG pathway analysis using KAAS [41].

## Supporting information

https://hungria.imd.ufrn.br/~danilo/files_paper/docs/FD_DaniloMartins_Supplementary.docx

## Data availability

The raw reads produced in this study were deposited in the NCBI database Sequence Read Archive (https://www.ncbi.nlm.nih.gov/Traces/sra) under the accession number **PRJNA665688**.

## Author information

## Contributions

AMRS, AV and TVS performed animal management and collected all samples. AMRS, AV and TVS. generated RNA-Seq data. JESS. and DLM. conceived the experimental design. JESS performed sequencing quality control. DLM performed transcriptome assembly, annotation and expression analyses. ACMFC and PAS contributed to these analyses. JESS and SS conceived and JESS supervised the study. DLM and ACMFC drafted the manuscript. All authors have read and approved the final manuscript.

## Corresponding author

Correspondence to Jorge E. S. de Souza

## Ethics declarations Competing interests

The authors declare that they have no competing interests.

