## Supplementary material for "Characterization and analysis of the transcriptome in *Arapaima gigas* using multi-tissue RNA-sequencing": https://hungria.imd.ufrn.br/~danilo/files_paper/docs/FD_DaniloMartins_Supplementary.docx

5 - Núcleo de Pesquisas em Oncologia, Universidade Federal do Pará, Belém, PA, Brazil.

Correspondence:
Dr. Jorge E. S. de Souza


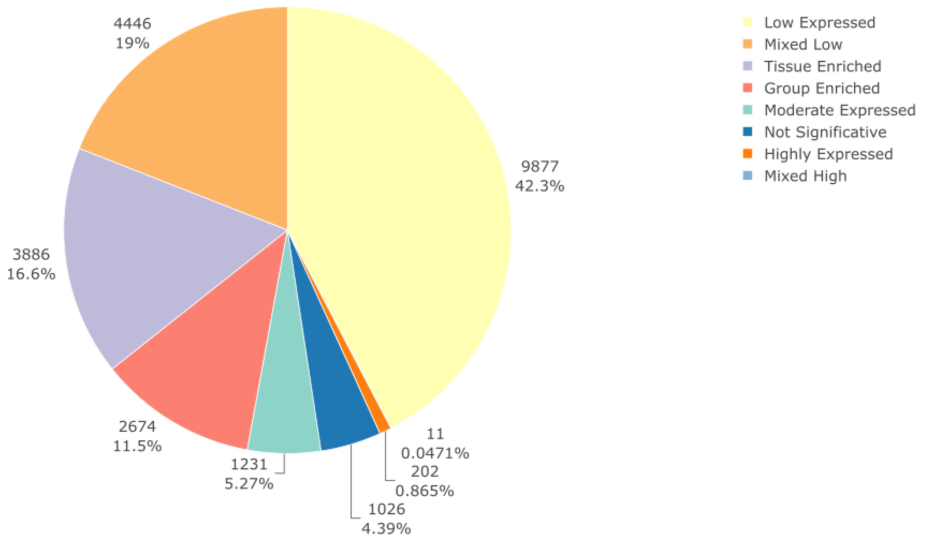


**Figure S1. Distribution of categorized genes according expression levels:** Highly Expressed, Tissue Enriched, Group Enriched, Low Expressed, Mixed Low, Mixed High, Moderate Expressed, Not Significative.

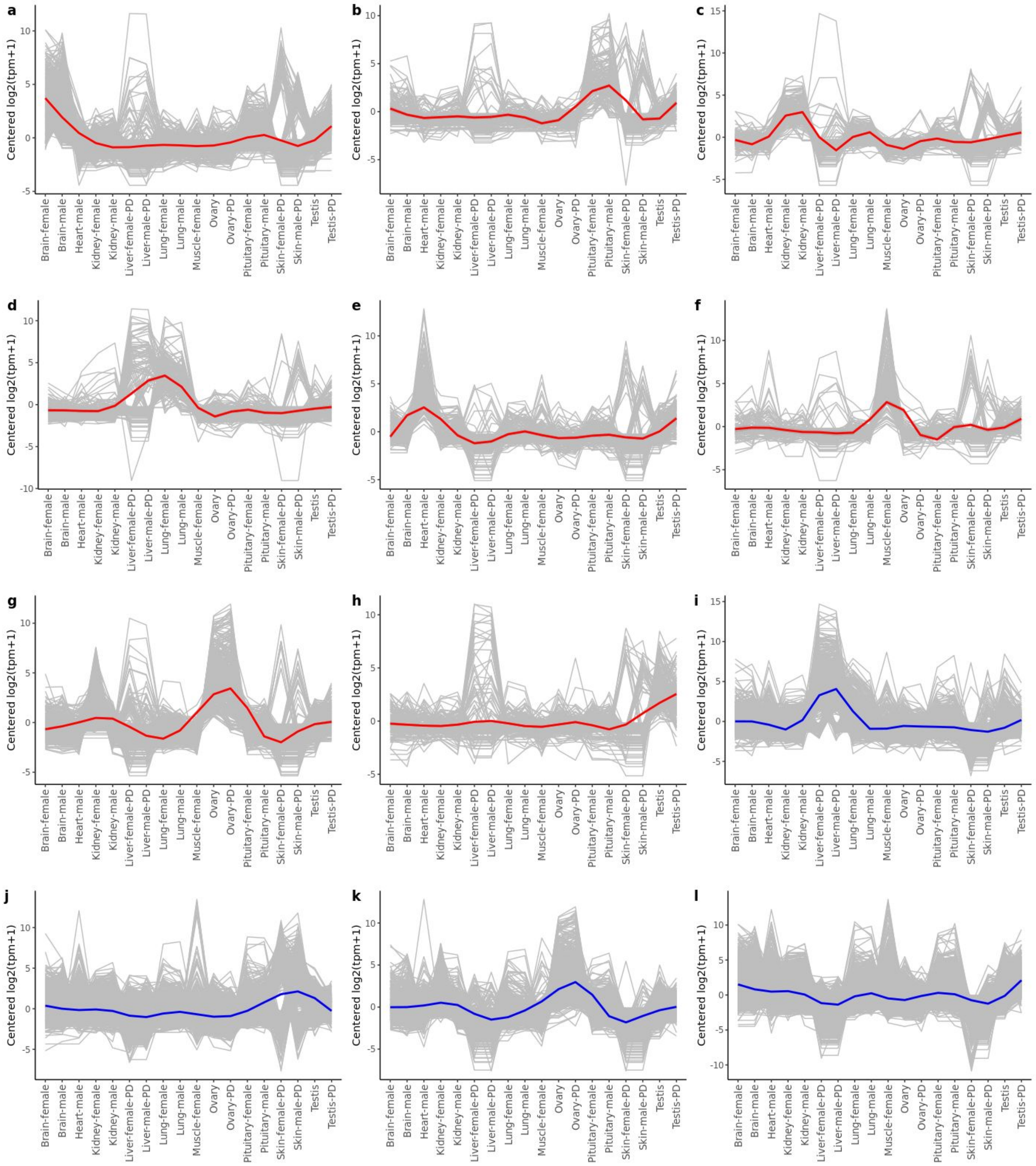


**Figure S2. Relative expression of tissue specific clusters across tissues.** The y-axis of each graph represents the mean-centered log2 (TPM+1) value. Expression levels of single genes is shown in gray, and the mean expression of the genes in **(a)** brain, **(b)** pituitary, **(c)** kidney, **(d)** lung, **(e)** heart, **(f)** muscle, **(g)** ovary, and **(h)** testis clusters is shown in blue. The mean expression of genes identified in public data is shown in red for (**i**) liver, (**j**) skin, (**k**) ovary, and (**l**) testis.

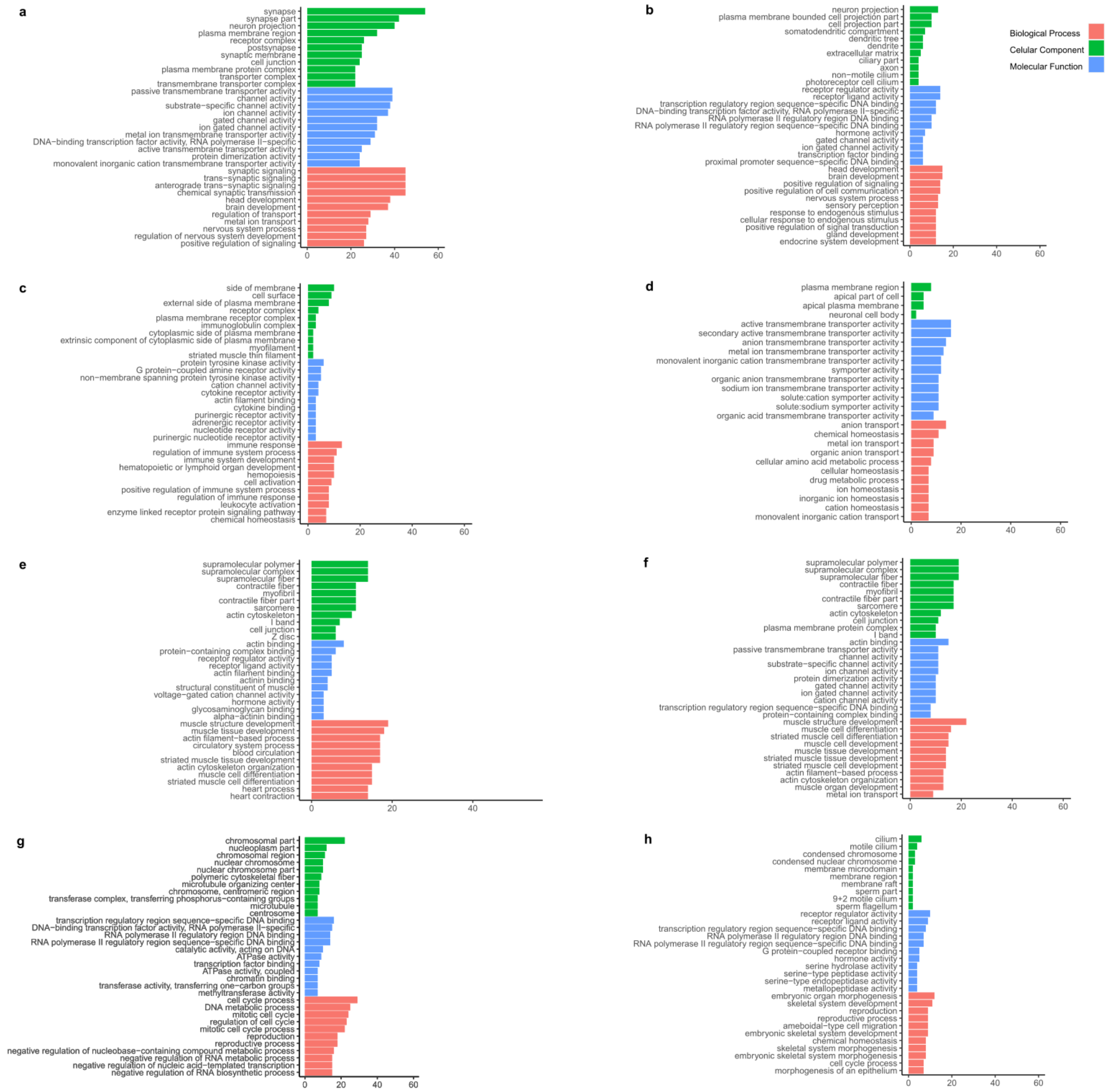


**Figure S3. Top 10 enriched terms in functional enrichment analysis for (a) brain, (b) pituitary, (c) kidney, (d) lung, (e) heart, (f) muscle, (g) ovary, and (h) testis.**

| RefSeq | Description | Family | Brain | Heart | Pituitary | Kidney | Lung | Muscle | Ovary | Testis |
|---|---|---|---|---|---|---|---|---|---|---|
| XP_018587820.1 | homeobox protein engrailed-1-B isoform X2 | Homeobox | 2.47428623561779 | -0.382159912426587 | -0.369063088317154 | -0.367642700596815 | -0.344420775075026 | -0.337856279356239 | -0.360645575023121 | -0.312497904822843 |
| XP_029113003.1 | T-cell leukemia homeobox protein 3-like | Homeobox | 2.47487196762474 | -0.354277166807348 | -0.352383060842596 | -0.354277166807348 | -0.354277166807348 | -0.351103072745402 | -0.354277166807348 | -0.354277166807348 |
| XP_018617074.1 | nucleoside diphosphate kinase A-like isoform X1 | TF_others | 2.45696427314429 | -0.431702071021161 | -0.0662771217363549 | -0.379219408822119 | -0.415728340540038 | -0.4216063084932 | -0.407104377071024 | -0.335326645460393 |
| XP_018601787.1 | zinc finger protein ZIC 2 | zf-C2H2 | 2.47486540555637 | -0.353950180329043 | -0.347696228726066 | -0.355638125490212 | -0.353821227861849 | -0.352762494769003 | -0.355359022889983 | -0.355638125490212 |
| XP_018609148.1 | zinc finger protein ZIC 1 | zf-C2H2 | 2.47444912377407 | -0.358696351412364 | -0.361663406690252 | -0.361663406690252 | -0.360911942384911 | -0.308187203215783 | -0.361663406690252 | -0.361663406690252 |
| XP_018584695.1 | zinc finger protein ZIC 2-like | zf-C2H2 | 2.47486990381373 | -0.35482882311614 | -0.35224243810755 | -0.35482882311614 | -0.354267899008423 | -0.35482882311614 | -0.35409640945978 | -0.349776687889552 |
| XP_029113608.1 | TSC22 domain family protein 1 isoform X2 | TSC22 | 2.44496090208784 | -0.122469369033453 | -0.100073626896212 | -0.366782463303483 | -0.461822436779219 | -0.457796078295266 | -0.438834179302084 | -0.497182748478127 |
| XP_018618301.2 | DNA-binding protein inhibitor ID-4 | bHLH | 2.46823240779954 | -0.382709966374414 | -0.194615347219016 | -0.355819350444212 | -0.309072732264086 | -0.416068711517105 | -0.415782595932201 | -0.394163704048507 |
| XP_018612338.2 | homeobox protein engrailed-2b-like | Homeobox | 2.4748662064435 | -0.355765020939559 | -0.352784583186711 | -0.351124146503211 | -0.355765020939559 | -0.349048667894774 | -0.354613746040131 | -0.355765020939559 |
| XP_029114226.1 | serine/threonine-protein phosphatase PP1-beta catalytic subunit-like | THAP | 2.45415801334903 | -0.176598144453574 | -0.159129705334506 | -0.411048739857572 | -0.499588062279597 | -0.423295193589236 | -0.479290656387072 | -0.305207511447475 |
| XP_029114962.1 | homeobox protein Nkx-2.3-like isoform X1 | Homeobox | -0.409025657564503 | 2.45181377316275 | -0.409025657564503 | -0.0167504940365311 | -0.408635505532485 | -0.40603588512984 | -0.400348282350055 | -0.401992290984832 |
| XP_018591281.1 | heart- and neural crest derivatives-expressed protein 2 | bHLH | -0.380473649425379 | 2.47102317924906 | -0.380473649425379 | -0.37865820265738 | -0.365234198015477 | -0.380473649425379 | -0.36854440384517 | -0.2171654264549 |
| XP_018609176.1 | influenza virus NS1A-binding protein homolog A-like | ZBTB | -0.0869692388251501 | 2.45204251129965 | -0.39526504553081 | -0.538964331097184 | -0.277344846566326 | -0.292600307660969 | -0.437465147423525 | -0.423433594195688 |
| XP_018604994.1 | uncharacterized protein LOC108932842 | ZBTB | -0.384064124426518 | 2.46897381060826 | -0.203113822086803 | -0.423707822251376 | -0.311008525661219 | -0.377345881732948 | -0.400966134029559 | -0.368767500419841 |
| XP_029109346.1 | myosin-7-like | MBD | -0.365424328294227 | 2.47385545436304 | -0.365424328294227 | -0.363567528350003 | -0.365424328294227 | -0.365424328294227 | -0.365424328294227 | -0.283166284541901 |
| XP_018600630.2 | myosin-7-like | MBD | -0.353878771670999 | 2.47487306491411 | -0.353788048984504 | -0.354111684776897 | -0.352054969140514 | -0.354165354470487 | -0.354014331424388 | -0.352859904446324 |
| XP_017334548.1 | PREDICTED: myosin-7 | MBD | -0.357757017191451 | 2.47474502051339 | -0.356570059250951 | -0.358308125440507 | -0.357184381357743 | -0.35845364483405 | -0.35787525234932 | -0.328596540089372 |
| XP_018585205.1 | desmin | Homeobox | -0.358188329443994 | 2.47484645692027 | -0.358159551909078 | -0.345198409329479 | -0.355652714484376 | -0.355793899830652 | -0.353974089689936 | -0.347879462232757 |
| KPP69492.1 | alpha-actinin-2-like | Homeobox | -0.347616824066907 | 2.4748602712963 | -0.354467994601568 | -0.350540327848063 | -0.356183022187192 | -0.356615460326921 | -0.357167199140108 | -0.352269443125541 |
| XP_018587267.1 | serine/threonine-protein phosphatase PP1-gamma catalytic subunit-like | THAP | -0.338943429670889 | 2.47373225936658 | -0.373999777065811 | -0.363560502789431 | -0.367510247691869 | -0.373999777065811 | -0.37139649112861 | -0.284322033954156 |
| KPP67237.1 | pituitary-specific positive transcription factor 1-like, partial | Pou | -0.353833484097051 | -0.353833484097051 | 2.47487325864685 | -0.352041659019439 | -0.353664179142155 | -0.353833484097051 | -0.353833484097051 | -0.353833484097051 |
| XP_029105685.1 | achaete-scute homolog 1 | bHLH | -0.346284996905716 | -0.354932070577451 | 2.47486277392713 | -0.354581549258273 | -0.354267945453338 | -0.354932070577451 | -0.354932070577451 | -0.354932070577451 |
| XP_018588679.2 | cytoplasmic polyadenylation element-binding protein 2 isoform X1 | TF_others | -0.374573319463396 | -0.273143405685892 | 2.46942203683832 | -0.428343940839045 | -0.416621880557163 | -0.238874328156803 | -0.387728136950466 | -0.35013702518556 |
| KPP72215.1 | TSC22 domain family protein 3-like | TSC22 | -0.391006866664488 | -0.416806780581548 | 2.45803881957804 | -0.199586508689759 | -0.443754769336989 | -0.529965610805957 | -0.231797446595257 | -0.245120836904045 |
| XP_018586815.1 | LIM domain transcription factor LMO4 | Homeobox | -0.303134565703242 | -0.29755712081831 | 2.45703447990469 | -0.259575885630132 | -0.194024553509162 | -0.538753334778576 | -0.515755846599031 | -0.348233172866239 |
| XP_029114458.1 | homeobox protein SIX6 | Homeobox | -0.345075741676593 | -0.355122829081758 | 2.47485878498046 | -0.355122829081758 | -0.355122829081758 | -0.355122829081758 | -0.354168897895079 | -0.355122829081758 |
| KPP60961.1 | TSC22 domain family protein 3-like | TSC22 | -0.185589463985023 | -0.493048738453413 | 2.44627274301581 | -0.368302421559611 | -0.496299286532125 | -0.397902548938461 | -0.0650221287388956 | -0.440108154808276 |
| XP_018599168.1 | insulinoma-associated protein 1a | zf-C2H2 | -0.322182327801153 | -0.358673069594843 | 2.47467103865877 | -0.358308816372356 | -0.358665953630935 | -0.359210591280397 | -0.359032832398725 | -0.358597447580362 |
| AAL69332.1 | x-box binding protein 1A | TF_bZIP | -0.344946617386944 | -0.304253368634862 | 2.46512989370472 | -0.214083972623648 | -0.426568952268121 | -0.513675641513857 | -0.307752074771316 | -0.353849266505974 |
| XP_029114505.1 | insulinoma-associated protein 1b-like | zf-C2H2 | -0.16510505142747 | -0.38662711504945 | 2.46759351366939 | -0.385355480235811 | -0.372832369290067 | -0.38662711504945 | -0.384419267567694 | -0.38662711504945 |
| XP_018599564.1 | homeobox protein Nkx-2.6 | Homeobox | -0.36789428841467 | -0.36789428841467 | -0.36789428841467 | 2.47394939852523 | -0.365693655682428 | -0.36789428841467 | -0.348154399614187 | -0.288524189569941 |
| XP_018613401.1 | interferon regulatory factor 4-like | IRF | -0.355348406972323 | -0.412683870228532 | -0.16408214800438 | 2.45872564645506 | -0.216217601499149 | -0.48425780634929 | -0.478866507694119 | -0.347269305707269 |
| XP_018595497.1 | pleckstrin-like | TF_others | -0.458681988036805 | -0.405076055116382 | -0.168139061938312 | 2.4520135028168 | -0.212252026394249 | -0.492965561474011 | -0.501110707305306 | -0.213788102551735 |
| XP_029107298.1 | T-cell leukemia homeobox protein 1-like | Homeobox | -0.353553390593274 | -0.353553390593274 | -0.353553390593274 | 2.47487373415292 | -0.353553390593274 | -0.353553390593274 | -0.353553390593274 | -0.353553390593274 |
| XP_018597381.1 | bile acid receptor-like | THR-like | -0.372579205370203 | -0.382070973727122 | -0.380487847094353 | 2.4707317740135 | -0.212371835199607 | -0.38026111323452 | -0.382070973727122 | -0.360889825660572 |
| XP_018618877.1 | interferon regulatory factor 8 isoform X2 | IRF | -0.362621080858956 | -0.362621080858956 | -0.328201427491007 | 2.47461497299658 | -0.333308141210799 | -0.362621080858956 | -0.362621080858956 | -0.362621080858956 |
| XP_018606396.2 | T-cell leukemia homeobox protein 1-like isoform X1 | Homeobox | -0.358511005006917 | -0.323623270555368 | -0.358511005006917 | 2.47468930059687 | -0.358511005006917 | -0.358511005006917 | -0.358511005006917 | -0.358511005006917 |
| XP_029110909.1 | tripartite motif-containing protein 16-like | zf-C2H2 | -0.536569548102443 | -0.0532312884609849 | -0.192987302207485 | 2.44021709364238 | -0.278255051580389 | -0.478943700676161 | -0.501766994122166 | -0.398463208492756 |
| XP_018595312.1 | myosin-11-like isoform X2 | MBD | -0.356189979050358 | -0.430718855336198 | -0.194415804722606 | 2.45595516889202 | -0.240734411390955 | -0.523611465202908 | -0.484993206743655 | -0.225291446445338 |
| XP_018618648.1 | neuronal acetylcholine receptor subunit alpha-3-like isoform X1 | Homeobox | -0.384819642816481 | -0.285460044238368 | -0.377754443183959 | 2.4722283285162 | -0.275211651063377 | -0.384819642816481 | -0.379343261581054 | -0.384819642816481 |
| XP_018611882.2 | forkhead box protein I1 | Fork_head | -0.353553390593274 | -0.353553390593274 | -0.353553390593274 | -0.353553390593274 | 2.47487373415292 | -0.353553390593274 | -0.353553390593274 | -0.353553390593274 |
| XP_018602401.1 | ETS-related transcription factor Elf-3-like | ETS | -0.386087434988629 | -0.374303994377058 | -0.180714161933276 | -0.411669732997424 | 2.46758009094833 | -0.424541771341 | -0.370476248020432 | -0.319786747290511 |
| XP_023652245.1 | homeobox protein EMX1 | Homeobox | -0.343941623428141 | -0.357537647049618 | -0.357537647049618 | -0.357537647049618 | 2.47482440183727 | -0.357537647049618 | -0.343194543161041 | -0.357537647049618 |
| KPP73591.1 | hepatocyte nuclear factor 1-beta-like | Homeobox | -0.365771944026777 | -0.365771944026777 | -0.365771944026777 | -0.365771944026777 | 2.47373532466043 | -0.365771944026777 | -0.279103660499774 | -0.365771944026777 |
| XP_029104705.1 | CCAAT/enhancer-binding protein beta-like | TF_bZIP | -0.245736435351722 | -0.392635198446374 | -0.536391127861337 | -0.141882617927904 | 2.42564203122734 | -0.558692575183847 | -0.530407929892485 | -0.0198961465636738 |
| KPP73182.1 | hypothetical protein Z043_107753 | Fork_head | -0.326553410964572 | -0.106553317980089 | -0.398308752056735 | -0.407524757433674 | 2.46151451073609 | -0.407524757433674 | -0.407524757433674 | -0.407524757433674 |
| KPP73964.1 | Motor neuron and pancreas homeobox protein 1-like | Homeobox | -0.355373438273763 | -0.355373438273763 | -0.355373438273763 | -0.350166332196184 | 2.47486298639555 | -0.34782946283055 | -0.355373438273763 | -0.355373438273763 |
| XP_018586251.1 | hepatocyte nuclear factor 1-beta-A-like isoform X2 | Homeobox | -0.371947545674705 | -0.351734044896989 | -0.375797110148225 | -0.375797110148225 | 2.47239955615987 | -0.375797110148225 | -0.245529524995276 | -0.375797110148225 |
| XP_018596631.1 | CCAAT/enhancer-binding protein gamma-like | TF_bZIP | -0.420501876514281 | -0.420501876514281 | -0.39255163923081 | 0.0346909387557501 | 2.44416745610665 | -0.420501876514281 | -0.404299249574466 | -0.420501876514281 |
| XP_018598424.2 | keratin, type I cytoskeletal 18-like isoform X1 | Homeobox | -0.410041699107175 | -0.31664812475813 | -0.342954478466713 | -0.400881642962492 | 2.47122873634221 | -0.422754866867483 | -0.305868603252166 | -0.272079320928054 |
| XP_018590727.1 | calpain-3-like isoform X1 | TF_others | -0.345294329359459 | -0.365662314875758 | -0.363584271913912 | -0.351564926811572 | -0.338289994327818 | 2.47473623759856 | -0.344678085434278 | -0.365662314875758 |
| TRY87356.1 | hypothetical protein DNTS_017220 | zf-C2HC | -0.342729945203946 | -0.289708744021716 | -0.37594404304067 | -0.379550830769947 | -0.409561619929552 | 2.47320059391806 | -0.322662705625626 | -0.353042705326603 |
| XP_018603312.2 | LOW QUALITY PROTEIN: kelch-like protein 31 | ZBTB | -0.417985599992532 | 0.0416999400187011 | -0.391750487086202 | -0.423438056988087 | -0.420704295376645 | 2.44303228298171 | -0.425988054086398 | -0.404865729470544 |
| XP_015193830.1 | PREDICTED: myoblast determination protein 1 | bHLH | -0.353553390593274 | -0.353553390593274 | -0.353553390593274 | -0.353553390593274 | -0.353553390593274 | 2.47487373415292 | -0.353553390593274 | -0.353553390593274 |
| XP_029107917.1 | myosin heavy chain, fast skeletal muscle-like | MBD | -0.353593435651008 | -0.353754474425364 | -0.353754474425364 | -0.353754474425364 | -0.353705143781644 | 2.47487352572101 | -0.353754474425364 | -0.352557048586904 |
| XP_030632386.1 | myosin heavy chain, fast skeletal muscle isoform X1 | MBD | -0.354006541306626 | -0.353515367416965 | -0.352444406572177 | -0.354006541306626 | -0.354006541306626 | 2.47487329286896 | -0.354006541306626 | -0.352887353653317 |
| XP_029107693.1 | myosin heavy chain, fast skeletal muscle-like | MBD | -0.354430484398168 | -0.354430484398168 | -0.352477295020812 | -0.354430484398168 | -0.354430484398168 | 2.47487129984307 | -0.354148691633422 | -0.350523375596165 |
| XP_018613590.2 | calsequestrin-1-like | zf-C2H2 | -0.35394945235449 | -0.353855803718753 | -0.353993266698084 | -0.353718472602584 | -0.353975862424506 | 2.47487316528594 | -0.353425033804637 | -0.351955273682886 |
| XP_018599939.2 | uncharacterized protein LOC108929717 isoform X2 | Homeobox | -0.285526133466708 | -0.309321692052435 | -0.344778232271508 | -0.319351001255658 | -0.408398638346135 | 2.47042966353315 | -0.471089257139876 | -0.331964709000833 |
| XP_018619088.2 | calsequestrin-1-like isoform X1 | zf-C2H2 | -0.350132240971893 | -0.354402194761423 | -0.354402194761423 | -0.353280355277342 | -0.354402194761423 | 2.47487112695763 | -0.353849751662697 | -0.354402194761423 |
| XP_018593551.1 | diencephalon/mesencephalon homeobox protein 1-like | Homeobox | -0.378141220573574 | -0.384692264829195 | -0.380532646484721 | -0.172229381345124 | -0.384168667277733 | -0.383692529694888 | 2.46814897503443 | -0.384692264829195 |
| XP_018608842.1 | LIM/homeobox protein Lhx8 isoform X1 | Homeobox | -0.401422908995561 | -0.396199827892723 | -0.389436052928027 | -0.175530172750559 | -0.387780113918085 | -0.406590339935204 | 2.4672719857528 | -0.310312569332638 |
| XP_030056874.1 | uncharacterized protein LOC115468892 isoform X1 | Fork_head | -0.410351831154714 | -0.410351831154714 | -0.410351831154714 | 0.014099896108103 | -0.409776410512085 | -0.410351831154714 | 2.44743567017755 | -0.410351831154714 |
| XP_018615926.2 | iroquois-class homeodomain protein irx-1-A | Homeobox | -0.377236412970488 | -0.386521403041259 | -0.372137042123231 | -0.19756277509552 | -0.382602420653137 | -0.386565369900235 | 2.46984058113343 | -0.367215157349556 |
| XP_018617970.1 | factor in the germline alpha isoform X1 | bHLH | -0.388707565751099 | -0.388707565751099 | -0.388707565751099 | -0.181880795277288 | -0.362661863615901 | -0.388707565751099 | 2.46872453825697 | -0.369351616359383 |
| XP_026871912.1 | P43 5S RNA-binding protein-like | zf-C2H2 | -0.386857060152697 | -0.370208680103678 | -0.38960967145959 | -0.175433442531194 | -0.390118755528049 | -0.388710339123553 | 2.46829407925162 | -0.367356130352856 |
| XP_018595085.1 | lipopolysaccharide-induced tumor necrosis factor-alpha factor homolog isoform X1 | zf-LITAF-like | -0.376624892950727 | -0.375943621375009 | -0.376330390250929 | -0.211998227652209 | -0.376624892950727 | -0.376624892950727 | 2.47077181108106 | -0.376624892950727 |
| XP_018599866.1 | mRNA decay activator protein ZFP36L2-like | zf-CCCH | -0.397655353709848 | -0.398486364338509 | -0.392415669739238 | -0.075438555712308 | -0.398154719898482 | -0.398486364338509 | 2.4591233920754 | -0.398486364338509 |
| XP_029623646.1 | P43 5S RNA-binding protein-like isoform X2 | zf-C2H2 | -0.391455313131195 | -0.391455313131195 | -0.391198961121023 | -0.142824085137555 | -0.385106562909867 | -0.391455313131195 | 2.46575869676642 | -0.372263148204389 |
| XP_018617215.1 | GATA-type zinc finger protein 1 isoform X1 | zf-GATA | -0.398304410301753 | -0.399089294025417 | -0.399089294025417 | -0.0640707590599035 | -0.399089294025417 | -0.399089294025417 | 2.45782163948874 | -0.399089294025417 |
| XP_029114101.1 | protein odd-skipped-related 1 | zf-C2H2 | -0.375810988559269 | -0.249349143565915 | -0.467320411106675 | -0.431547854812046 | -0.464863035072854 | -0.171081892918525 | -0.30092972615977 | 2.46090305219505 |
| KPP65756.1 | Dmrt1a-like | DM | -0.366099233218929 | -0.366099233218929 | -0.336516431431225 | -0.366099233218929 | -0.335870604788618 | -0.366099233218929 | -0.337828156232085 | 2.47461212532764 |
| XP_018618734.1 | zinc finger protein basonuclin-1-like isoform X1 | zf-C2H2 | -0.310801837579973 | -0.375304561601629 | -0.358994693560729 | -0.368435086544463 | -0.369611116668716 | -0.375304561601629 | -0.315586603172608 | 2.47403846072975 |
| XP_029112307.1 | Wilms tumor protein isoform X2 | zf-C2H2 | -0.441712147842958 | -0.441712147842958 | -0.441712147842958 | 0.0148979287088178 | -0.385257639760068 | -0.441712147842958 | -0.307456612807523 | 2.4446649152306 |
| XP_018615968.1 | homeobox protein Hox-C6a-like | Homeobox | -0.384190691605655 | -0.384190691605655 | -0.384190691605655 | -0.384190691605655 | -0.223197746843884 | -0.335409452225007 | -0.375688768057993 | 2.4710587335495 |
| XP_018604651.1 | zinc finger and BTB domain-containing protein 7C | ZBTB | -0.370635524426485 | -0.301389390933786 | -0.367073882953544 | -0.379082510814138 | -0.294154468430382 | -0.430478678229314 | -0.329606925824839 | 2.47242138161249 |
| XP_029108653.1 | NACHT, LRR and PYD domains-containing protein 3-like | zf-C2H2 | -0.156085230128246 | -0.435012909369451 | -0.265936495818666 | -0.435012909369451 | -0.378896090735834 | -0.435012909369451 | -0.357007145139708 | 2.46296368993081 |
| RXM91507.1 | High mobility group-T protein | HMG | -0.357806545743192 | -0.341579028440823 | -0.357806545743192 | -0.357806545743192 | -0.349587283382224 | -0.357806545743192 | -0.35244010098476 | 2.47483259578057 |
| XP_018613546.1 | putative leucine-rich repeat-containing protein DDB_G0290503 | zf-C2H2 | -0.373990340609935 | -0.344137504497291 | -0.374243022876808 | -0.338643038245373 | -0.365792807997496 | -0.373754680135009 | -0.303567446597214 | 2.47412884095913 |
| XP_028828455.1 | keratin, type I cytoskeletal 19-like | Homeobox | -0.399112667421626 | -0.378539512272902 | -0.395136287550069 | -0.396625058954898 | -0.256620908146823 | -0.336423599769304 | -0.309243077158128 | 2.47170111127375 |

**Figure S4. Top 10 highly expressed transcription factors in each tissue-specific cluster along with their scaled mean expression values.**

| Refseq | Symbol | Family | Brain−female | Pituitary−female | Kidney−female | Lung−female | Muscle−female | Ovary | Brain−male | Heart−male | Pituitary−male | Kidney−male | Lung−male | Testis |
|---|---|---|---|---|---|---|---|---|---|---|---|---|---|---|
| XP_018582739.1 | znf706 | TF_others | −0.516500573007686 | −0.489694730707608 | 0.540509210697038 | 0.139387787010994 | −0.554727600441325 | 3.00588015368287 | −0.243866313147676 | −0.257273700074177 | −0.561643027053987 | −0.309373566665691 | −0.473087023325838 | −0.279610616966912 |
| XP_029105180.1 | LOC108928113 | bHLH | −0.505151041108581 | −0.536123534570778 | −0.251451621371452 | 2.45078460974793 | −0.85044378714497 | 1.3771889554207 | −0.267403400679025 | 0.517980789239425 | −0.582394381308757 | −0.719635578502826 | 0.113721287269238 | −0.747072296990906 |
| XP_029109103.1 | gata6 | zf−GATA | −0.481097764182357 | −0.507208609494243 | −0.38772058806771 | −0.443794162348523 | −0.507208609494243 | −0.292768619192652 | −0.491524229154157 | 2.18067504042334 | −0.507208609494243 | −0.406589082051323 | −0.239785505871989 | 2.0842307389281 |
| XP_029102676.1 | nme4 | TF_others | −0.440763395353487 | −0.343023170131149 | 0.375843313316652 | 0.104119055155917 | −0.440763395353487 | 3.06952727379703 | −0.40532976104928 | −0.440763395353487 | −0.440763395353487 | −0.40506523227837 | −0.351316109014015 | −0.281701788382834 |
| XP_029108941.1 | klf4 | zf−C2H2 | −0.431973230283103 | −0.474477764586962 | −0.26097818540102 | −0.437793314105048 | −0.126151966651527 | −0.41567834234141 | −0.491736798560535 | 0.842503158077411 | −0.392976190565717 | −0.278100642730196 | −0.485688435339982 | 2.95305171248809 |
| XP_018595294.1 | tfa | TF_others | −0.431053401391746 | −0.432717210751233 | −0.43264960913162 | 0.666779451334368 | −0.431978248526926 | −0.43140372596117 | −0.431593971146109 | −0.391988880894688 | −0.432495722726034 | −0.426417509365656 | 2.97910424595845 | 0.196414582602359 |
| XP_017537984.1 | twist1b | bHLH | −0.40924179676745 | −0.282203256893113 | 0.257935828263637 | −0.396120108718339 | −0.40924179676745 | 3.11881293521795 | −0.354468371235418 | −0.146974523975172 | −0.364083038726851 | −0.363184118000014 | −0.400474313313754 | −0.250757439084023 |
| XP_018584355.2 | neurod1 | bHLH | −0.395302259230563 | 0.378386775886697 | −0.427082279478447 | −0.427082279478447 | −0.425855937009084 | −0.426915278588781 | 3.0114660494423 | −0.42657727374783 | 0.418720935298164 | −0.426744020771773 | −0.425932152843791 | −0.427082279478447 |
| XP_029108219.1 | dmrta1 | DM | −0.389344164914301 | 0.613522175442432 | −0.402851661678973 | −0.404708528298707 | −0.353389511045113 | −0.443906494783772 | −0.218033508318089 | −0.2064864186501 | 3.03761146925907 | −0.394193929414923 | −0.415070142955317 | −0.423149284642212 |
| TKS82811.1 | | Homeobox | −0.385356731937021 | −0.385356731937021 | 0.813090336038657 | −0.385356731937021 | −0.385356731937021 | 2.98261884354036 | −0.376290897785467 | −0.385356731937021 | −0.385356731937021 | −0.385356731937021 | −0.385356731937021 | −0.33656442629738 |
| XP_030056874.1 | LOC115468892 | Fork_head | −0.368308831765634 | −0.368308831765634 | 0.643574598187759 | −0.368308831765634 | −0.368308831765634 | 3.03814192951903 | −0.368308831765634 | −0.368308831765634 | −0.368308831765634 | −0.368308831765634 | −0.366937041816083 | −0.368308831765634 |
| XP_018621066.1 | cpeb1 | TF_others | −0.364919363890262 | −0.364919363890262 | 0.59780599142161 | −0.364919363890262 | −0.364919363890262 | 3.05138764748101 | −0.364919363890262 | −0.364919363890262 | −0.364919363890262 | −0.364919363890262 | −0.364919363890262 | −0.364919363890262 |
| XP_018613000.1 | LOC108937492 | zf−C2H2 | −0.356101511872025 | −0.466975000313797 | −0.149812716425132 | −0.245056470684192 | −0.465628149108152 | −0.352683907191177 | −0.39329317719383 | 0.216430661747251 | −0.497600298454386 | −0.0540761629601035 | −0.343482785238289 | 3.10827951769383 |
| XP_018608842.1 | lhx8a | Homeobox | −0.354851100144389 | −0.324917214342541 | 0.163396078280942 | −0.330612754226731 | −0.357679024244737 | 3.14170858149329 | −0.347922598968247 | −0.345026911882317 | −0.348664651062854 | −0.316048615098328 | −0.338936369320815 | −0.240445420483272 |
| KPP72215.1 | | TSC22 | −0.354001397373897 | 3.16790001130699 | −0.201186298683321 | −0.341032231577993 | −0.400400501045405 | −0.216334072053661 | −0.275234408419006 | −0.330544814596025 | −0.279566012897593 | −0.191712661276291 | −0.353328689818268 | −0.224558923565537 |
| XP_018602401.1 | LOC108931228 | ETS | −0.353007134319399 | −0.240786288858486 | −0.387730239125683 | 3.12545785101953 | −0.394085791802425 | −0.355250342233103 | −0.379920687700844 | −0.357999825479857 | −0.197100889928273 | −0.381949286500468 | 0.241212536119695 | −0.318839901190687 |
| XP_018617215.1 | zglp1 | zf−GATA | −0.352483269345275 | −0.354375122756706 | 0.453140613852294 | −0.354375122756706 | −0.354375122756706 | 3.08871876030333 | −0.354375122756706 | −0.354375122756706 | −0.354375122756706 | −0.354375122756706 | −0.354375122756706 | −0.354375122756706 |
| XP_018599866.1 | cth1 | zf−CCCH | −0.351827999631632 | −0.339180675607911 | 0.425925062007026 | −0.353833857379311 | −0.353833857379311 | 3.09495396316813 | −0.353833857379311 | −0.353833857379311 | −0.353833857379311 | −0.353833857379311 | −0.353033348280439 | −0.353833857379311 |
| XP_018588679.2 | cpeb2 | TF_others | −0.348085817896913 | 3.13181379890484 | −0.377348671358798 | −0.3717021239638 | −0.24307993072208 | −0.348561209729944 | −0.330392980887141 | −0.267363797943768 | 0.220350220338929 | −0.37733635905233 | −0.366369856083877 | −0.32192327160512 |
| XP_028828455.1 | LOC114785964 | Homeobox | −0.346063393092882 | −0.320323179076216 | −0.328319708427505 | −0.267076477050771 | −0.263495310703855 | −0.23030730014992 | −0.334016664351368 | −0.314919757858647 | −0.350046385377775 | −0.345685500697685 | −0.0650324925442277 | 3.16528616933085 |
| KPP79452.1 | | zf−LITAF−like | −0.344510120836186 | −0.33430367029758 | 0.317303627874314 | −0.344510120836186 | −0.344510120836186 | 3.11759112994894 | −0.344510120836186 | −0.344510120836186 | −0.344510120836186 | −0.344510120836186 | −0.344510120836186 | −0.344510120836186 |
| XP_029623646.1 | LOC115203203 | zf−C2H2 | −0.342827120333489 | −0.342827120333489 | 0.261180513103746 | −0.328328306515662 | −0.342827120333489 | 3.1277326445608 | −0.342827120333489 | −0.342827120333489 | −0.342204356361962 | −0.342827120333489 | −0.341902716290016 | −0.31951505649597 |
| XP_026871912.1 | si:dkey−208k4.2 | zf−C2H2 | −0.339833518332978 | −0.339833518332978 | 0.147626518210596 | −0.339833518332978 | −0.337511527398022 | 3.1451807825986 | −0.330671237241287 | −0.314957979641401 | −0.337382111168727 | −0.302679913257507 | −0.338623259526048 | −0.311480717577271 |
| XP_018617970.1 | figla | bHLH | −0.339451553553549 | −0.339451553553549 | 0.164652383786166 | −0.30555905543823 | −0.339451553553549 | 3.14279296201745 | −0.339451553553549 | −0.339451553553549 | −0.339451553553549 | −0.339451553553549 | −0.309862225950408 | −0.31586318954013 |
| XP_018609757.2 | rxrgb | RXR−like | −0.33664245226437 | −0.37359847681753 | 0.53647468802711 | −0.318478720286319 | −0.382305241763818 | 3.06724150396184 | −0.376584785083766 | −0.382305241763818 | −0.33643883229939 | −0.382305241763818 | −0.377013545247974 | −0.338043654698148 |
| XP_018615926.2 | irx7 | Homeobox | −0.334956256253701 | −0.299678679330168 | 0.116154188648457 | −0.327810041461813 | −0.336547648645591 | 3.14938274635283 | −0.315369102119471 | −0.336493991913072 | −0.338200232608884 | −0.327935562203438 | −0.33561256541273 | −0.312932835052417 |
| XP_023690222.1 | LOC111855444 | Homeobox | −0.329576428664688 | 3.15363519215835 | −0.329576428664688 | −0.282010462592993 | −0.320967109870344 | −0.326596766408067 | −0.329576428664688 | −0.329576428664688 | 0.0807083735436897 | −0.329576428664688 | −0.327310654842511 | −0.329576428664688 |
| XP_029622666.1 | LOC115202771 | bHLH | −0.327949395952044 | −0.327949395952044 | 0.131942015303383 | −0.327949395952044 | −0.327949395952044 | 3.14755194421706 | −0.327949395952044 | −0.327949395952044 | −0.327949395952044 | −0.327949395952044 | −0.327949395952044 | −0.327949395952044 |
| XP_018593551.1 | LOC108925792 | Homeobox | −0.325657600442691 | −0.329406920310993 | 0.179188525081056 | −0.335208750612406 | −0.333990588541458 | 3.14093495343696 | −0.328795205007312 | −0.335208750612406 | −0.330873717412726 | −0.331840435674801 | −0.333932759290815 | −0.335208750612406 |
| XP_018595085.1 | si:dkeyp−75b4.8 | zf−LITAF−like | −0.323439043839115 | −0.323439043839115 | 0.0786857622749591 | −0.323439043839115 | −0.323439043839115 | 3.15415325708732 | −0.323439043839115 | −0.322606990903463 | −0.322719677745892 | −0.323439043839115 | −0.323439043839115 | −0.323439043839115 |
| XP_023678976.1 | fev | ETS | −0.312183144694614 | 3.16598597896267 | −0.312183144694614 | −0.312183144694614 | −0.312183144694614 | −0.312183144694614 | −0.312183144694614 | −0.312183144694614 | −0.0441545320165282 | −0.312183144694614 | −0.312183144694614 | −0.312183144694614 |
| XP_029102934.1 | znf385b | zf−C2H2 | −0.311038819037851 | 0.539753962381475 | −0.452662014034666 | −0.452662014034666 | −0.394224755021765 | −0.452662014034666 | 0.351030456709307 | −0.452662014034666 | 2.9831132532115 | −0.452662014034666 | −0.452662014034666 | −0.452662014034666 |
| XP_018613546.1 | si:ch211−14a17.10 | zf−C2H2 | −0.308822661688606 | −0.309925704241964 | −0.260450227850038 | −0.299054929604257 | −0.309085937408401 | −0.223237183078653 | −0.309925704241964 | −0.272860009884214 | −0.309440792843198 | −0.271828798174705 | −0.299639989912116 | 3.17427193892812 |
| XP_018596829.2 | LOC108927773 | zf−C2H2 | −0.270645307426969 | −0.279761980765159 | −0.284616970989767 | −0.292827120814307 | −0.292827120814307 | −0.292827120814307 | 3.17534810087405 | −0.292827120814307 | −0.292827120814307 | −0.291162205654413 | −0.292198911151902 | −0.292827120814307 |
| XP_018601787.1 | zic2a | zf−C2H2 | −0.265562167809772 | −0.285787445461585 | −0.292481204987576 | −0.292481204987576 | −0.290705978396826 | −0.292308905271267 | 3.17533577972049 | −0.2914391778901 | −0.289369347809921 | −0.292481204987576 | −0.290237937130718 | −0.292481204987576 |
| XP_029110917.1 | LOC108932644 | bHLH | −0.262643525342389 | −0.292823316188244 | −0.292823316188244 | −0.292823316188244 | −0.291177068770825 | −0.292823316188244 | 3.1753025276592 | −0.292823316188244 | −0.292823316188244 | −0.281469485131801 | −0.290249235096483 | −0.292823316188244 |
| XP_018584695.1 | LOC108920431 | zf−C2H2 | −0.260761703342486 | −0.28900261583305 | −0.292200697264368 | −0.292200697264368 | −0.292200697264368 | −0.291747880203747 | 3.17530069269486 | −0.292200697264368 | −0.292200697264368 | −0.292200697264368 | −0.291507111055311 | −0.289077198674057 |
| XP_029113003.1 | LOC108925283 | Homeobox | −0.258699431272183 | −0.289745348764669 | −0.292089097061531 | −0.292089097061531 | −0.290125300502572 | −0.292089097061531 | 3.17528285703167 | −0.292089097061531 | −0.292089097061531 | −0.292089097061531 | −0.292089097061531 | −0.292089097061531 |
| XP_018609148.1 | zic1 | zf−C2H2 | −0.233978377238384 | −0.297695707347801 | −0.297695707347801 | −0.297695707347801 | −0.264357151420308 | −0.297695707347801 | 3.17481018222926 | −0.295845962379105 | −0.297695707347801 | −0.297695707347801 | −0.296758739756857 | −0.297695707347801 |
| XP_018587820.1 | LOC108922262 | Homeobox | −0.209997177617127 | −0.313143359962019 | −0.294895858854888 | −0.276630813675447 | −0.285299494246593 | −0.299622068814919 | 3.1741384861667 | −0.313143359962019 | −0.29668122417182 | −0.313143359962019 | −0.302219459175985 | −0.269362309723865 |
| XP_029106150.1 | LOC108935213 | zf−C2H2 | −0.209424732779488 | −0.304132242022787 | −0.304132242022787 | −0.304132242022787 | −0.264226015092608 | −0.30348871284822 | 3.17410891049283 | −0.304132242022787 | −0.304132242022787 | −0.304132242022787 | −0.303145398827905 | −0.269030598807881 |
| XP_029107436.1 | npas4l | bHLH | −0.198305917741496 | 0.0300910853060199 | −0.321096275258738 | −0.34954767692076 | −0.344019840619429 | −0.333737130752102 | 3.15644541699605 | −0.275078376448006 | −0.354027579770041 | −0.344190608708781 | −0.332524977739921 | −0.334008118342795 |
| XP_018618734.1 | bnc1 | zf−C2H2 | −0.159132926165576 | −0.312342757131423 | −0.300291339401761 | −0.303835939975868 | −0.315795928111217 | −0.242695798992163 | −0.314544708654999 | −0.315795928111217 | −0.27931962088818 | −0.314482811498216 | −0.313817343305468 | 3.17205510223609 |
| XP_029102751.1 | neurod2 | bHLH | −0.153632314963839 | −0.306589992072511 | −0.306589992072511 | −0.306589992072511 | −0.306589992072511 | −0.306589992072511 | 3.17228382279716 | −0.306589992072511 | −0.306589992072511 | −0.306589992072511 | −0.304105816794481 | −0.261825754458752 |
| XP_018620030.1 | LOC108941726 | bHLH | −0.133690161710021 | −0.303793728286764 | −0.303793728286764 | −0.303793728286764 | −0.303793728286764 | −0.303793728286764 | 3.17162744457786 | −0.303793728286764 | −0.303793728286764 | −0.303793728286764 | −0.303793728286764 | −0.303793728286764 |
| KPP69899.1 | | Homeobox | −0.0836132222032036 | −0.124368481940114 | −0.465323743695805 | −0.546685857231283 | −0.558047298818184 | −0.385798028407034 | −0.228798758514994 | 0.133029238302952 | −0.0140863456247807 | −0.303291917008671 | −0.516949343719662 | 3.09393375686078 |
| XP_030285865.1 | ppp3ca | THAP | −0.0672724665411007 | −0.259302050169486 | −0.339330161166864 | −0.352119174449673 | −0.0605991647594003 | −0.428369216314064 | 3.1519168510773 | −0.429823637571274 | −0.234258447526099 | −0.31789551028175 | −0.390292717511773 | −0.272654304785821 |
| XP_029114226.1 | LOC108922004 | THAP | 0.0733004282936767 | −0.229920157961277 | −0.403745577987972 | −0.432758937018905 | −0.370215680745608 | −0.408700725166441 | 3.14154990465149 | −0.200663605986415 | −0.147395335137576 | −0.319852086903134 | −0.41254286553436 | −0.289055360503475 |
| XP_018615057.2 | hopx | Homeobox | 0.173515120341294 | 0.245260427106681 | −0.136214382475539 | 0.535572155199342 | −0.780367773353045 | 2.72829838252534 | −0.670458329007566 | −0.82028584691987 | 0.467527065622457 | −0.754413123204527 | −0.217488222693959 | −0.770945473140602 |
| XP_018617295.1 | LOC108940023 | CTF/NFI | 0.376041456575275 | −0.0983066034672606 | −0.566057322166005 | −0.522230191774426 | −0.244431006035335 | −0.564678242996035 | 3.05447173625582 | −0.163809711216406 | −0.279225716792863 | −0.5294055696905 | −0.397639213860984 | −0.0647296148312789 |
| XP_029112959.1 | sox19b | HMG | 0.540767302492662 | −0.290174064268685 | 0.210543939246999 | −0.50908424560769 | −0.50908424560769 | 2.98486015522678 | −0.0498220345656104 | −0.50908424560769 | −0.356661986774939 | −0.50908424560769 | −0.494092083318755 | −0.50908424560769 |
| XP_018580904.2 | her2 | bHLH | 0.825186261784145 | −0.00462444822252992 | 0.266203707742914 | −0.370016715559537 | −0.685471476775924 | 2.827704500222 | −0.521016493410328 | −0.687802245029136 | −0.11703759250963 | −0.253990295652974 | −0.591332957559868 | −0.687802245029136 |
| XP_018601601.1 | LOC108930709 | TF_bZIP | 1.40274554425872 | 2.66587853909792 | −0.467998959283766 | −0.512378482741915 | −0.508754196011862 | −0.51290744867322 | 0.0508665125791184 | −0.43283896673023 | −0.462870862463266 | −0.393984195485455 | −0.513717995737877 | −0.314039488808166 |
| XP_018617074.1 | LOC108939902 | TF_others | 3.0572510009277 | −0.0358285051756282 | −0.37880928681825 | −0.414020582989648 | −0.424627497155112 | −0.413523836987527 | 0.501550914758745 | −0.432357496213837 | −0.269298341991508 | −0.405537149341506 | −0.426233271165126 | −0.358565947848303 |
| XP_030628685.1 | nkx6.2 | Homeobox | 3.11949247142557 | −0.290763417983496 | −0.348657616282705 | −0.353455936218688 | −0.353455936218688 | −0.316230083083074 | 0.30413418331227 | −0.353455936218688 | −0.347239920076434 | −0.353455936218688 | −0.353455936218688 | −0.353455936218688 |

**Figure S5. Top 10 highly expressed male- and female-biased transcription factors.**

**Table S1. Top 10 ranked differentially expressed genes between males and females of *A. gigas.***

| Refseq | Symbol | Description | TPM.Female | TPM.Male | log2FC |
|---|---|---|---|---|---|
| XP_018590256.1 | *LOC108923784* | *GTP-binding protein Di-Ras2-like* | 466.060 | 0 | -Inf |
| XP_018589388.2 | *LOC108923234* | *retinoid-binding protein 7-like* | 444.691 | 0 | -Inf |
| KPP79776.1 | | *tubulin-specific chaperone A-like* | 391.899 | 0 | -Inf |
| XP_029107738.1 | *fam113* | *PC-esterase domain-containing protein 1A* | 167.912 | 0 | -Inf |
| PWS23252.1 | | *hypothetical protein DKP78_14105* | 111.593 | 0 | -Inf |
| AKA58770.1 | | *ferritin* | 103.732 | 0 | -Inf |
| XP_018598293.2 | *btg4* | *maternal B9.15 protein-like* | 79.007 | 0 | -Inf |
| XP_018613617.1 | *zp3f.2* | *zona pellucida sperm-binding protein 3-like* | 69.878 | 0 | -Inf |
| XP_018617215.1 | *zglp1* | *GATA-type zinc finger protein 1 isoform X1* | 59.001 | 0 | -Inf |
| XP_018621066.1 | *cpeb1* | *cytoplasmic polyadenylation element-binding protein 1 isoform X1* | 55.875 | 0 | -Inf |
| XP_029113759.1 | *LOC114912197* | *uncharacterized protein LOC114912197 isoform X2* | 0 | 37.514 | Inf |
| XP_018597916.1 | *LOC108928463* | *centrosomal protein of 164 kDa-like* | 0.079 | 21.261 | 8.080 |
| XP_018613905.1 | *slc1a3a* | *excitatory amino acid transporter 1-like* | 0.130 | 32.444 | 7.959 |
| XP_018601513.1 | *LOC108930641* | *cerebellin-1-like* | 0.3946 | 87.403 | 7.794 |
| KPP77175.1 | | *cerebellin-3 precursor-like* | 2.184 | 392.873 | 7.491 |
| XP_018589496.1 | *LOC108923312* | *cadherin-20-like isoform X1* | 0.371 | 55.960 | 7.237 |
| XP_018616543.1 | *dkk1b* | *dickkopf-related protein 1* | 0.310 | 40.921 | 7.045 |
| XP_018586657.1 | *LOC108921611* | *cortexin-1-like* | 0.230 | 26.658 | 6.856 |
| XP_029110917.1 | *LOC108932644* | *neurogenic differentiation factor 6-A-like* | 0.349 | 40.284 | 6.850 |
| XP_018582867.2 | *si:dkey-1h6.8* | *uncharacterized protein LOC108919389* | 0.327 | 35.812 | 6.775 |

**Table S2. KEGG pathway enrichment analysis with p-value ≤ 0.05 of female- and male-biased genes.**

| ID | Description | p-value | Count | Gender |
|---|---|---|---|---|
| ko04114 | Oocyte meiosis | 8.986821e-05 | 8 | Female |
| ko04110 | Cell cycle | 6.230763-04 | 7 | Female |
| ko00480 | Glutathione metabolism | 4.928937e-04 | 5 | Female |
| ko04914 | Progesterone-mediated oocyte maturation | 4.225671e-03 | 5 | Female |
| ko04115 | p53 signaling pathway | 1.045054e-02 | 4 | Female |
| ko00983 | Drug metabolism | 1.894772e-02 | 4 | Female |
| ko03010 | Ribosome | 4.625235e-02 | 4 | Female |
| ko00030 | Pentose phosphate pathway | 3.929582e-03 | 3 | Female |
| ko04020 | Calcium signaling pathway | 2.398148e-02 | 4 | Male |
| ko04144 | Endocytosis | 2.780246e-02 | 4 | Male |
| ko04371 | Apelin signaling pathway | 2.827757e-02 | 3 | Male |
| ko00590 | Arachidonic acid metabolism | 1.831551e-02 | 2 | Male |
